# Generative Molecular Design and Experimental Validation of Selective Histamine H1 Inhibitors

**DOI:** 10.1101/2023.02.14.528391

**Authors:** Kevin S. McLoughlin, Da Shi, Jeffrey E. Mast, John Bucci, John P. Williams, W. Derek Jones, Derrick Miyao, Luke Nam, Heather L. Osswald, Lev Zegelman, Jonathan Allen, Brian J. Bennion, Amanda K. Paulson, Ruben Abagyan, Martha S. Head, James M. Brase

## Abstract

Generative molecular design (GMD) is an increasingly popular strategy for drug discovery, using machine learning models to propose, evaluate and optimize chemical structures against a set of target design criteria. We present the ATOM-GMD platform, a scalable multiprocessing framework to optimize many parameters simultaneously over large populations of proposed molecules. ATOM-GMD uses a junction tree variational autoencoder mapping structures to latent vectors, along with a genetic algorithm operating on latent vector elements, to search a diverse molecular space for compounds that meet the design criteria. We used the ATOM-GMD framework in a lead optimization case study to develop potent and selective histamine H1 receptor antagonists. We synthesized 103 of the top scoring compounds and measured their properties experimentally. Six of the tested compounds bind H1 with *K_i_*’s between 10 and 100 nM and are at least 100-fold selective relative to muscarinic M2 receptors, validating the effectiveness of our GMD approach.

## INTRODUCTION

Automated design and optimization of compounds with desired pharmacological properties have been long-time goals of drug discovery. Advances in machine learningbased prediction of chemical properties and new approaches to learning-based generative design supported by high-performance computing are enabling new approaches to these goals. In recent years, a large body of work has developed incorporating both mechanistic and data-driven computational methods into these workflows^1–6^,^6–8^. These advances have enabled the development of generative model approaches to *de novo* drug design, here referred to as generative molecular design (GMD), in which new molecular structures are proposed by sampling a learned molecular generative model^9,10^. These techniques provide new tools for rapid *in silico* drug discovery considering all or most aspects of lead optimization, while maximizing the predictive value of existing discovery data and knowledge prior to lengthy synthesis and test cycles. Generally the GMD approach consists of three components: (1) a generative model, such as a generative adversarial network (GAN) or variational autoencoder (VAE) that structures the chemical space and maps it to a continuous numerical (i.e. latent space) representation, (2) a series of predictive models that relate chemical structures to the functional properties of molecules, and (3) an optimization framework that can navigate through the numerical chemical space representation toward structures for which the models predict more desirable properties. In some approaches, the generative model itself is optimized so that molecules sampled from the model tend to have the desired properties, for example by weighting molecules in the training data by Pareto efficiency^11^; by using joint entanglement of model weights^12^; or reinforcement learning^13^. Other approaches have applied techniques such as particle swarm optimization^14^ to iteratively sample molecules from the latent space and find regions corresponding to desired property ranges.

Most published examples of GMD workflows have involved *in silico* optimization of simple molecular properties, such as the octanol-water partition coefficient (log P)^9,15^ or inhibition of single proteins^14,16^. More recently, a few groups have experimentally validated molecules that were proposed by generative design techniques; these include compound classes such as DDR1 kinase inhibitors^17^, liver X receptor agonists^18^, selective JAK3 inhibitors^12^, and ligands for RXR and PPAR receptors^19^.

Here we report the development of the ATOM Generative Molecular Design (ATOM-GMD) platform, a scalable multiprocessing software framework to perform molecular optimization of many parameters simultaneously over large populations of proposed molecules. ATOM-GMD addresses one of the main challenges of lead optimization, which is to search chemical space to create molecular structures that jointly satisfy multiple objectives, including efficacy, safety, pharmacokinetic, and developability properties. It performs this search using a unique genetic algorithm approach, applying mutation and crossover operations directly to a working population of latent vectors, that samples a highly diverse molecular space while rewarding compounds that meet the objectives. We demonstrate the use of the ATOM GMD framework in a lead optimization case study to develop potent and selective histamine H1 receptor antagonists suitable for oral administration. We validated our approach by synthesizing 103 of the top scoring compounds and measuring their properties experimentally.

## RESULTS AND DISCUSSION

### Compounds Proposed by Generative Model

We generated candidate compound structures using two runs of the ATOM GMD software. The first run, initialized with a small set of 1,000 compounds, performed 200 generations of our genetic algorithm in 24 hours on a small compute cluster. It generated 199,000 latent vectors representing about 43,000 unique molecular structures. The second GMD run, initialized with 24,741 compound structures, produced 6 million latent vectors representing 1.7 million unique structures across 253 generations. The much larger initial set and additional generations allowed the GMD process to explore a broader range of chemical space. Figure 1 shows the number of compounds in each generation of the second run that were not previously proposed in the same run. As the GMD loop proceeds through generations, the optimizer “rediscovers” greater numbers of structures that were previously proposed. However, even in the last generations, over 4,000 out of 14,000 molecules were newly proposed in each generation.

**Figure 1.**
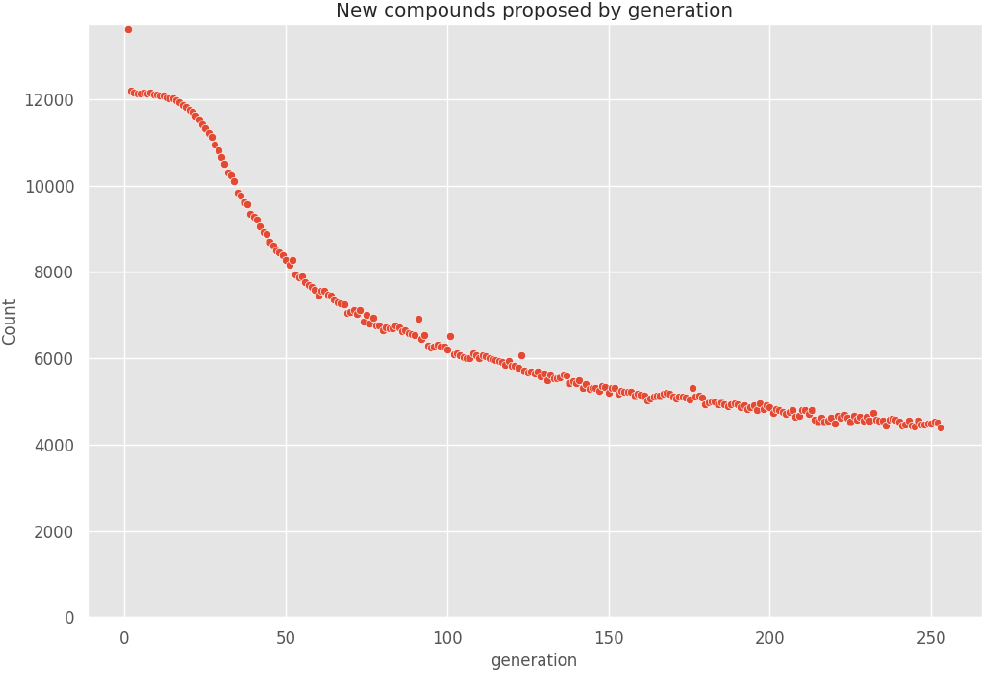
Numbers of unique molecular structures proposed in each generation of the second GMD run.

### Performance of the Multi-Parameter Optimization Process

A primary goal of this study was to design compounds predicted to bind strongly to histamine H1 receptors while binding weakly or not at all to muscarinic acetylcholine M2 receptors. The evolution of the working population of molecular structures toward a set satisfying these design criteria is illustrated in Figure 2, which plots the predicted H1 vs M2 *pK_i_* values for random samples of molecules in generations 10, 100 and 253 of the second run, compared with corresponding values for a sample of the initial compound set. The target *pK_i_* ranges (> 9 for H1 and < 6 for M2) are indicated by dashed lines. While the overall population maintains a wide range of values for each parameter, the median values increase for H1 and decrease for M2 in later generations, and the number of compounds in the target range steadily increases. As the majority of initial compounds already satisfied the M2 target criterion, the change in *pK_i_* distribution is most notable for H1.

**Figure 2.**
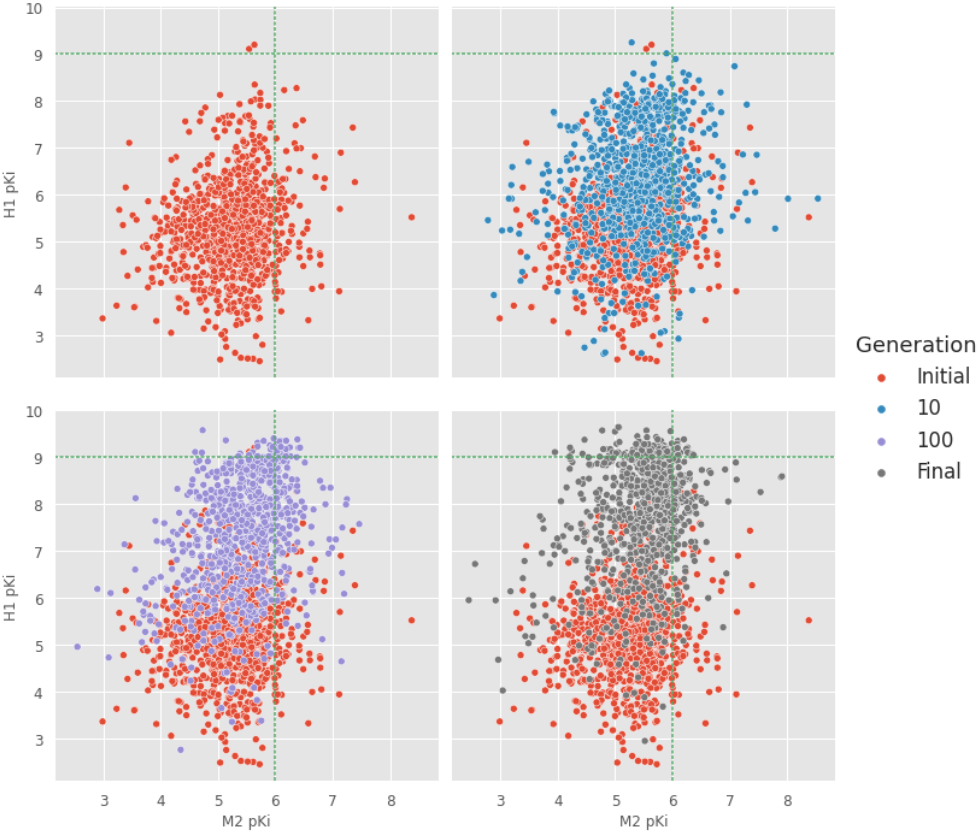
Comparison of the predicted H1 vs M2 *pK_i_* values of the initial library and the working population of molecules after 10, 100 and 253 generations in the second GMD run. To reduce overplotting, the plotted points for each generation shown are a random sample comprising 4% of the working population. The dashed horizontal and vertical lines represent the target minimum H1 and maximum M2 *pK_i_* values, respectively.

In addition to the H1 and M2 binding affinities, the GMD optimizer attempted to select for compounds satisfying design criteria for 10 other parameters, including applicability domain (AD) constraints on the property prediction models. The property optimization plots in Figure 3 show the changing distributions of individual parameter values over GMD generations. In each plot, the plotted error bars indicate the mean +/− one standard deviation range for the property, the X axis represents the generation number, the horizontal green line shows the target threshold for the property, and the arrows at the ends of the line are directed toward more desirable values. For properties with both maximum and minimum criteria, a green box is drawn spanning the desired range. A random sample of individual values is plotted for every fifth generation. The plots show that the target criteria were easily satisfied for some parameters, but harder to reach for others. In particular, the compounds meeting the H1 binding and hERG inhibition targets were generally in the tails of their respective *pK_i_* distributions. Nevertheless, the second GMD run generated 202 compounds that met or exceeded all 12 optimization criteria, including the AD constraints.

**Figure 3.**
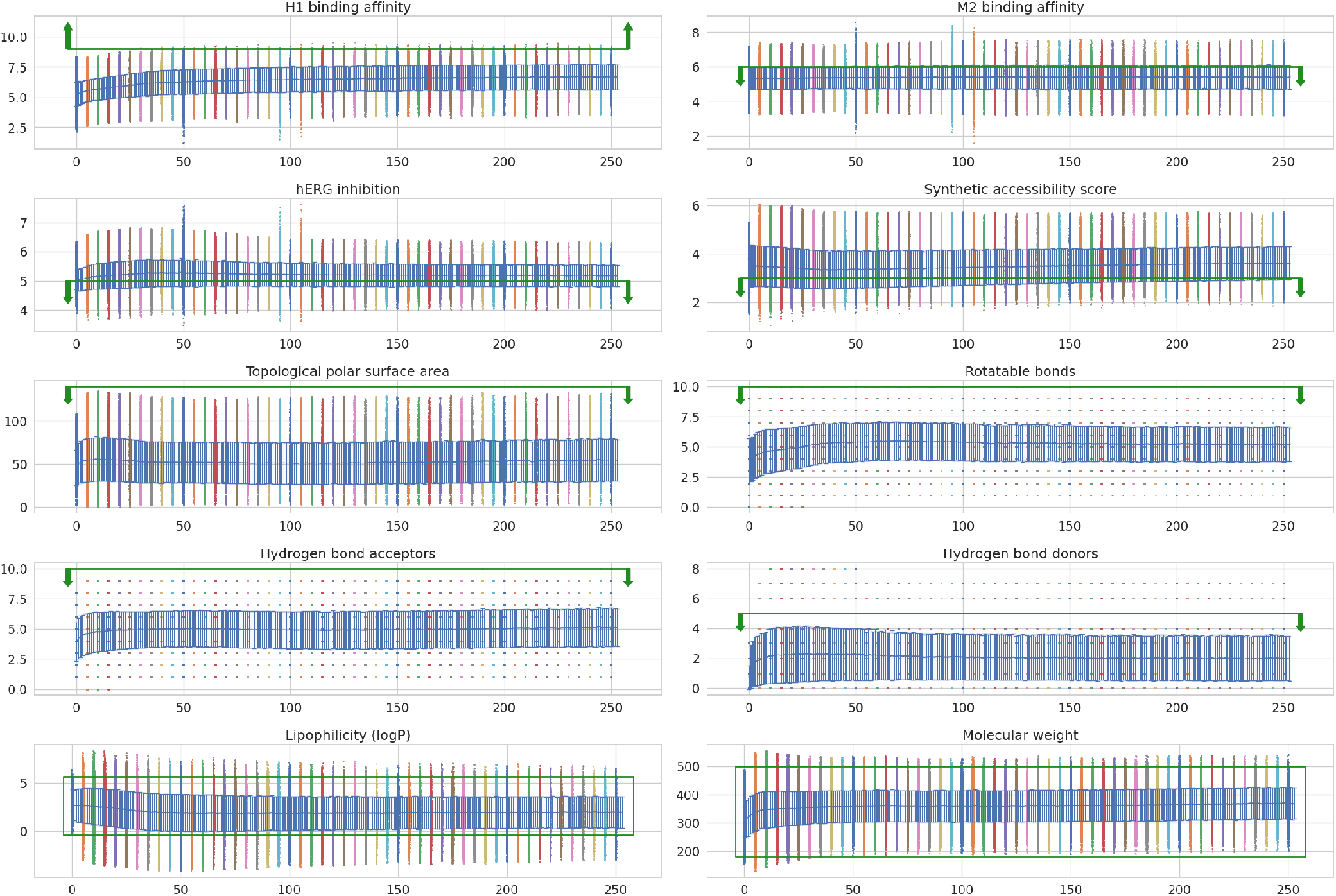
Predicted parameter value ranges by generation for the second GMD run

### Diversity of Generated Compounds

The ability to propose unique and novel compounds is an important characteristic for a generative molecular design platform. Ideally the GMD process should explore as wide a range of chemical structures as possible, while also exploiting newly gained knowledge about families of molecules with desirable properties. We performed several analyses to determine whether the ATOM GMD optimization algorithm maintained a diverse working population of molecular structures, not restricted to a narrow range of chemotypes identified in early generations. As a first step, we computed the average pairwise Tanimoto similarity between ECFP4 fingerprints of compounds in the working population of each generation in each GMD run; the results for the second run are plotted in Figure 4. The average similarity grows from about 0.12 to 0.2 during the first 50 generations, but then reaches a plateau. In our experience, compounds with a Tanimoto similarity below 0.2 usually share few major structural elements in common, suggesting that, at an aggregate level, the compound population generated by the GMD process is reasonably diverse.

**Figure 4.**
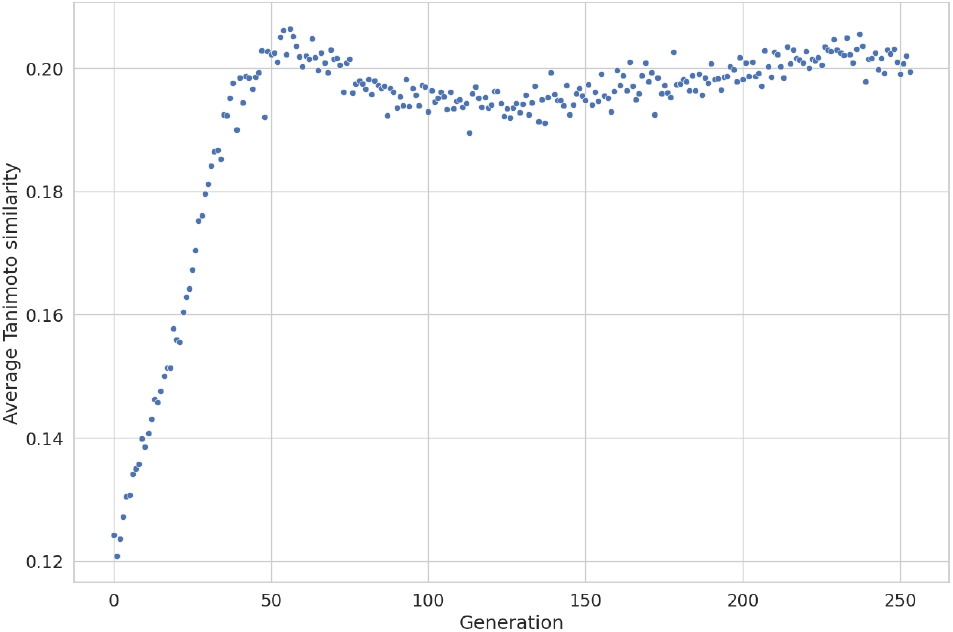
Average Tanimoto similarity between pairs of ECFP4 fingerprints in working population as a function of GMD loop generation.

To obtain a more granular view of the diversity of this population and its evolution, we performed Butina clustering^20^ of the full set of compounds from the first GMD run, using a similarity cutoff of 0.5, and selected the 50 largest clusters. We then used a stacked horizontal bar plot to visualize the changes in cluster representation over the course of the GMD run, as shown in Figure 5.

**Figure 5.**
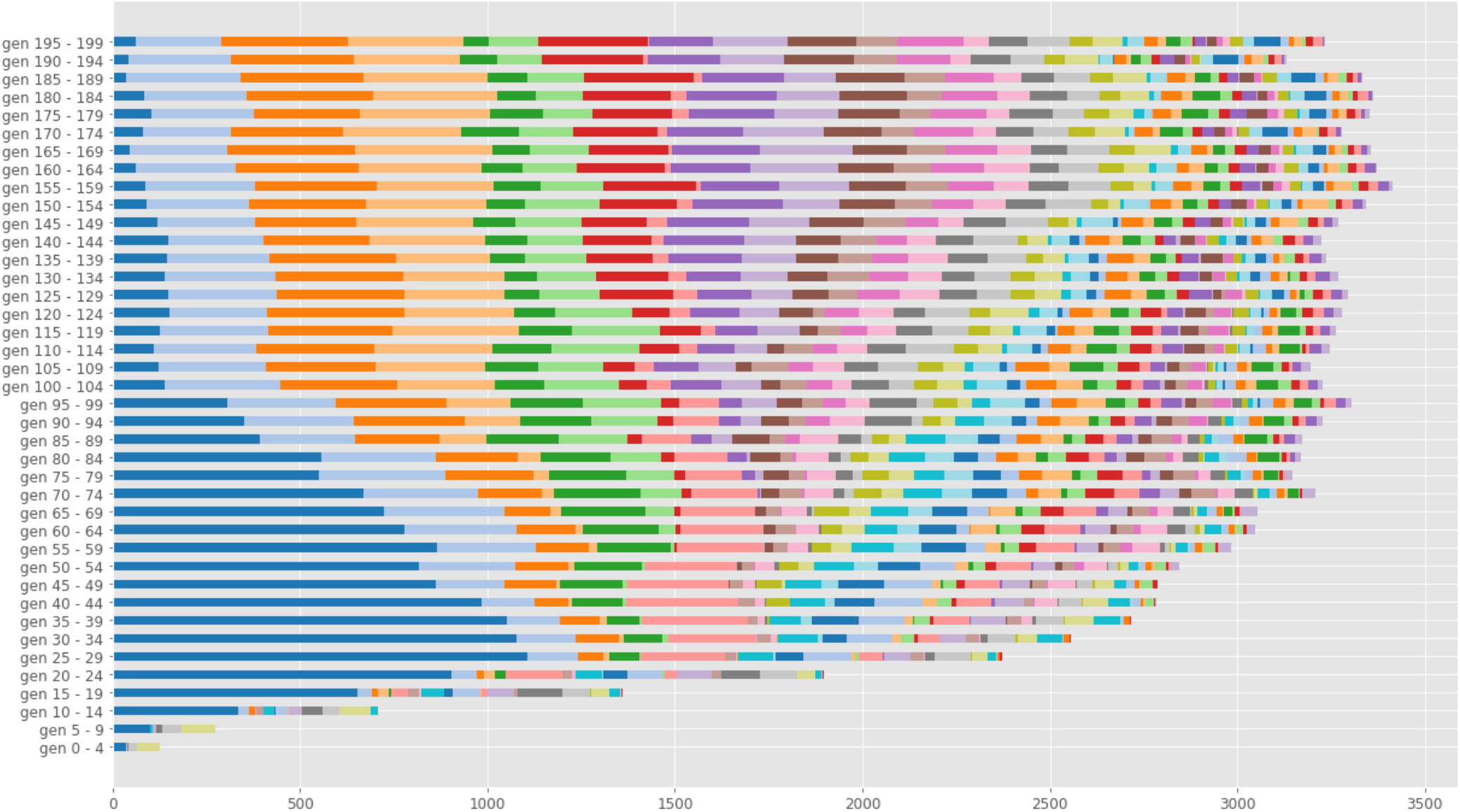
Evolution of Butina cluster representation over the course of the first GMD run.

The length of each colored bar segment corresponds to the number of cluster members in the population in a block of 5 generations. Thus, we see that the leftmost “dark blue” cluster grows rapidly over the first 30 generations, then shrinks as other clusters are born and grow to replace it in subsequent generations. In turn, some of the clusters that arise in later generations are seen to grow, shrink and sometimes disappear as they themselves are replaced. Thus, it appears that the genetic optimizer in our GMD platform creates a population of molecules that diversifies over time, rather than becoming dominated by a small set of chemotypes.

Yet another method to visualize the changes in chemical diversity over GMD generations is through a nonlinear 2D projection method such as t-distributed stochastic network embedding (t-SNE). When applied to ECFP4 fingerprint bit vectors, t-SNE groups together points representing structurally similar compounds. In Figure 6, we show t-SNE projections of compound fingerprints from the second GMD run initial set and the populations at generations 100, 200 and 250. A reference coordinate system was established by simultaneously projecting the fingerprints of 20,000 diverse compounds randomly selected from the ChEMBL database; these points are plotted in light blue. To reduce overplotting and create a more balanced representation, only 4,000 randomly selected points are plotted for each subset (the reference and initial sets and each selected generation). The initial compound set (dark blue points) is seen to cover most of the coordinate space spanned by the reference set, showing that the initial set is reasonably diverse; a few small clusters representing groups of related structures appear at the fringes of the plot area. The compounds from later generations are projected mainly into the lower right corner of the diagram and overlap with only a small set of initial and reference compounds, suggesting a great deal of structural novelty. Even at generation 250, the projections cover a broad area, indicating a strong degree of structural diversity.

**Figure 6.**
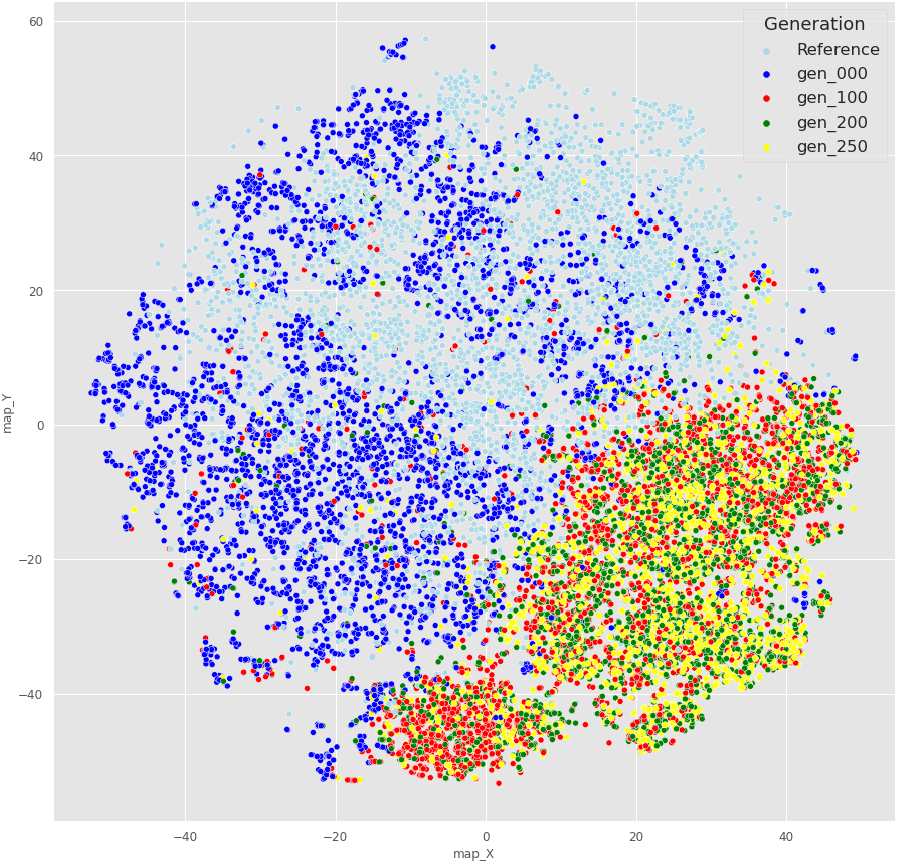
t-SNE projection of ECFP4 fingerprints from compounds in selected generations from the second GMD run.

### Selection and Synthesis of Proposed Molecules

From the first GMD run, the 2,000 top scoring compounds were selected as candidates for synthesis and testing. For the second run, we clustered the top 2,000 compounds by Tanimoto similarity using Butina’s algorithm and selected the best scoring representative of each cluster, yielding 566 additional candidates. Neurocrine chemists selected for synthesis 48 candidates from three chemical series (phthalazinone, benzodiazol and aminedimethylamino), and manually designed 7 additional members of the phthalazinone series. In addition, we found matches or near-matches to 95 candidates in the Enamine REAL virtual library and ordered these from Enamine; these included three exact matches that were also synthesized at Neurocrine. Details of the selection process are provided in the experimental section.

Of the 48 GMD compounds selected by Neurocrine chemists, 20 were successfully synthesized. Enamine was able to synthesize 79 of the 95 compounds requested. In addition, Neurocrine was able to produce all 7 of the phthalazinone series molecules that were designed manually. Thus, a total of 106 preparations (including the 3 compounds synthesized both by Neurocrine and by Enamine) were synthesized and tested for binding to H1 and M2 receptors and to the hERG channel. Table 1 summarizes the compound selection and synthesis process.

**Table 1.** Compound selection and synthesis results.

| Origin | Proposed compounds | Selected for synthesis | Successfully synthesized and tested |
| --- | --- | --- | --- |
| **GMD first run** |  |  |  |
| GMD compounds | 2000 | 45 | 20 |
| Enamine matches | 38 | 38 | 32 |
| **GMD second run** |  |  |  |
| GMD compounds | 566 | 3 | 0 |
| Enamine matches | 57 | 57 | 47 |
| **Manually designed at Neurocrine** | 7 | 7 | 7 |
| Total | 2668 | 150 | 106 |

### Binding of Compounds to H1 and M2 Receptors

Of the 106 compounds tested, 52 were measured as binding more than 50% of H1 receptors at 10 μM, while only 5 had greater than 50% binding to M2. An eight-point dilution series was performed for 40 of the 52 H1 binders, and binding measurements were taken for H1, M2 and hERG at each concentration. For those compounds that showed concentration-dependent binding, *IC_50_* and *K_i_* values were calculated. The detailed measurements are given together with the associated predictions in Supplementary File 2.

In Figure 7 we show the measured *pK_i_* values for H1 and M2 plotted against the values predicted by our QSAR models. In both plots, the dashed green line represents the identity (predicted *pK_i_* = measured *pK_i_*); the points clustered at the bottom of each plot represent compounds for which the *K_i_* was not measurable or was greater than 10 μM. Random jitter was added to the *Y* values for these points to reduce overplotting. Distinctive plot symbols are used for the 3 pairs of duplicate measurements from compounds synthesized twice. We see that our H1 binding model tended to overpredict *pK_i_*’s, with only one measurement exceeding the predicted value. Nevertheless, 8 compounds predicted to have p*K_i_* > 7 had measured *pK_i_*’s above this threshold; two of these were highly potent, with *K_i_* < 20 nM.

**Figure 7.**
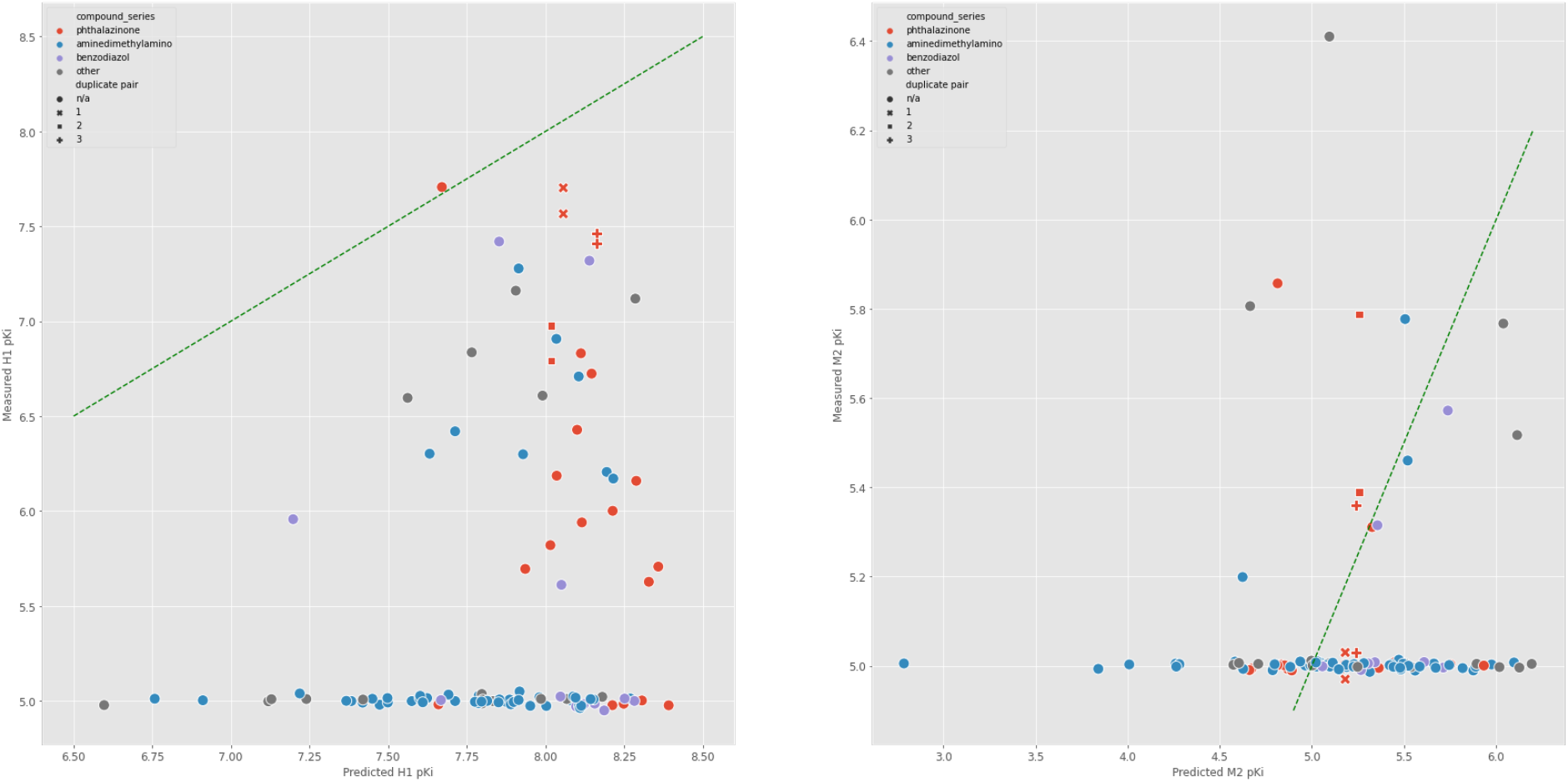
Measured vs predicted *pK_i_* values for binding to H1 and M2 receptors.

Our predictive model for M2 binding was somewhat less biased, with measured *pK_i_* values falling on both sides of the identity line. Most compounds had M2 *K_i_*’s that were greater than 10 μM or were not measurable, exceeding our design target (1 μM). All compounds with unmeasurable H1 *K_i_*’s also had unmeasurable M2 *K_i_*’s. Most of the compounds that bound strongly to H1 were also selective relative to M2. Figure 8 compares the measured *pK_i_* values for binding to H1 and M2; points above the upper dashed line represent compounds with greater than 100-fold selectivity for H1. Compounds for which M2 binding was below our measurement threshold are plotted along the vertical line at *pK_i_* = 5.0, with random jitter added in the *X* direction to separate overlapping points.

**Figure 8.**
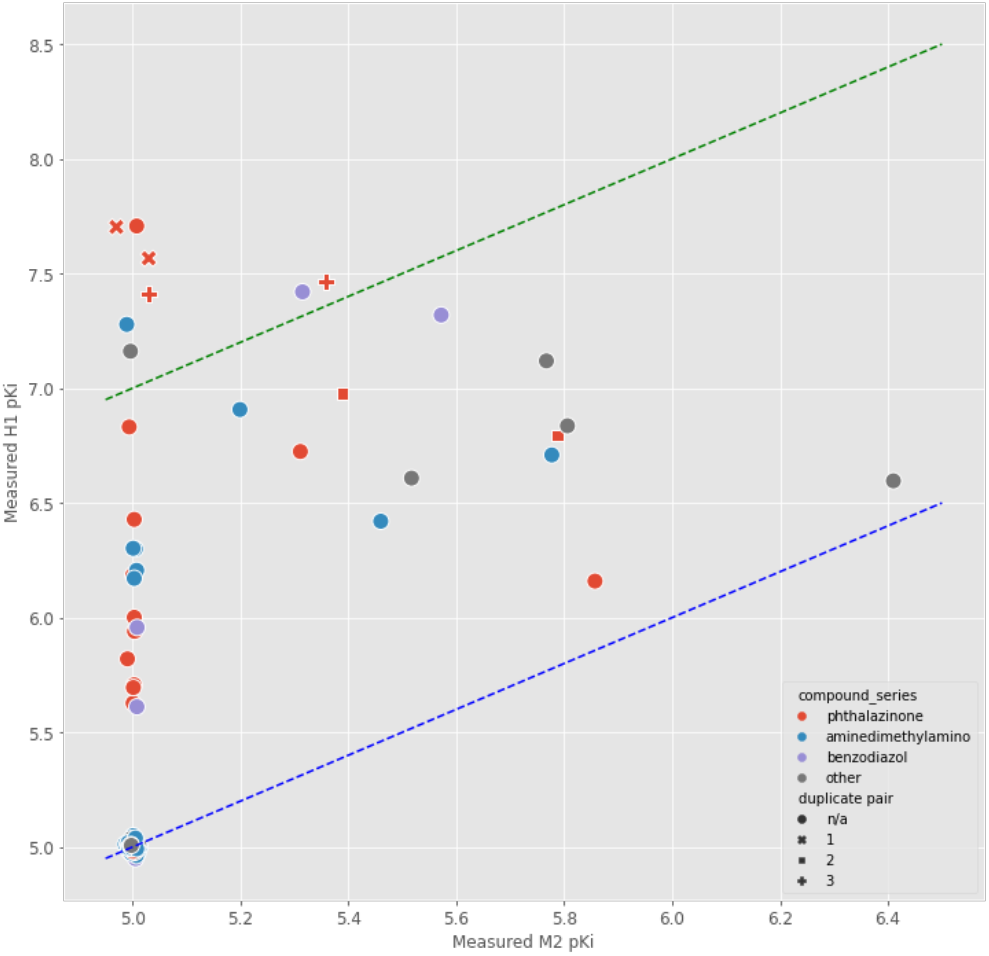
Measured *pK_i_* values for binding to H1 vs M2 receptors.

Structures, *K_i_*’s and selectivity ratios for the top 9 compounds ranked by H1 *K_i_* are shown in Table 2. All three of the compounds synthesized both at Enamine and at Neurocrine Biosciences Inc. (NBI) are included in this set; the measurement values are given separately for each batch. Measurement values for H1 and M2 *K_i_’s* were reasonably consistent between batches; the differences might be explained by variations in compound purity or by the intrinsic variability of binding measurements.

**Table 2.**
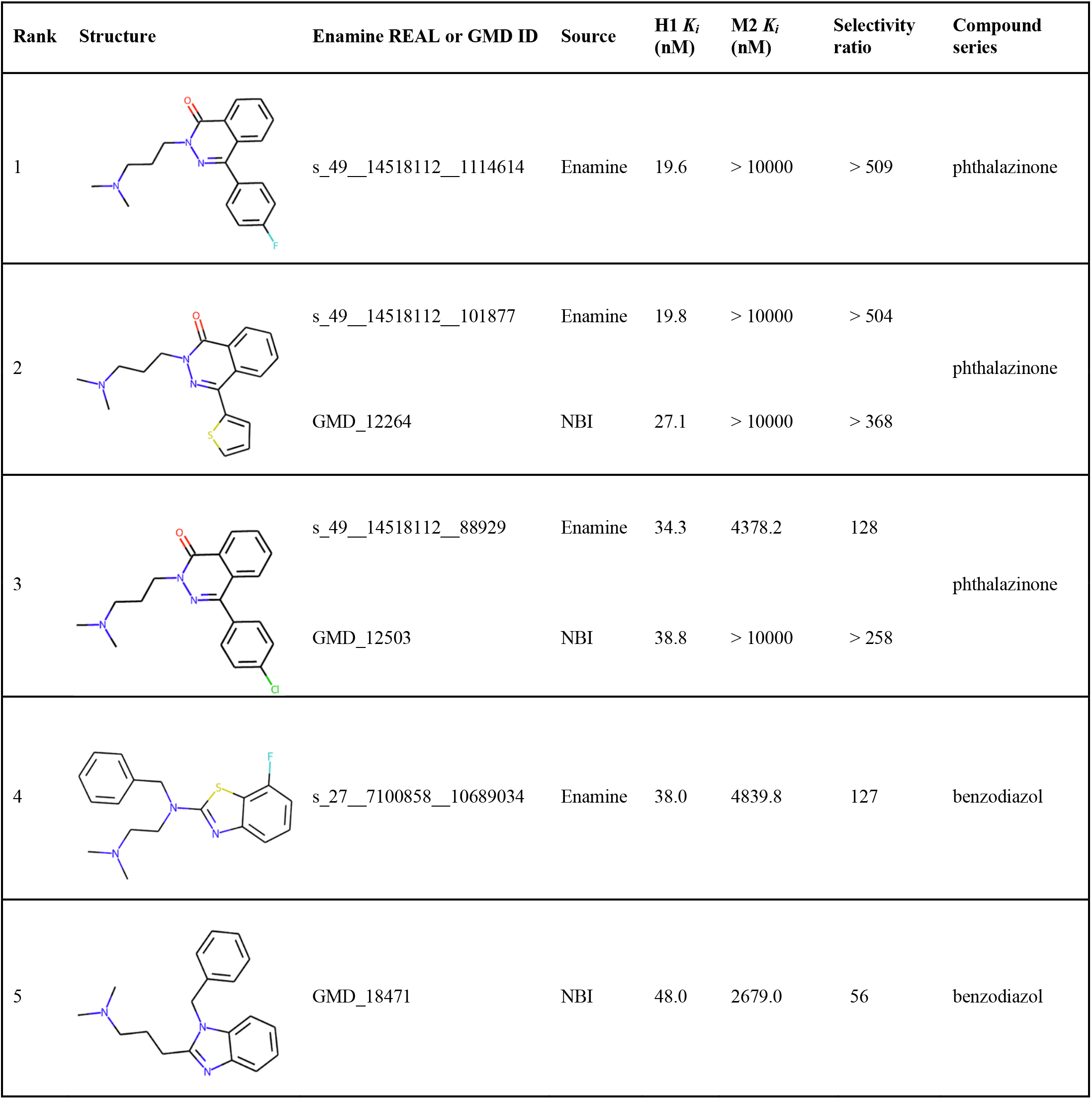
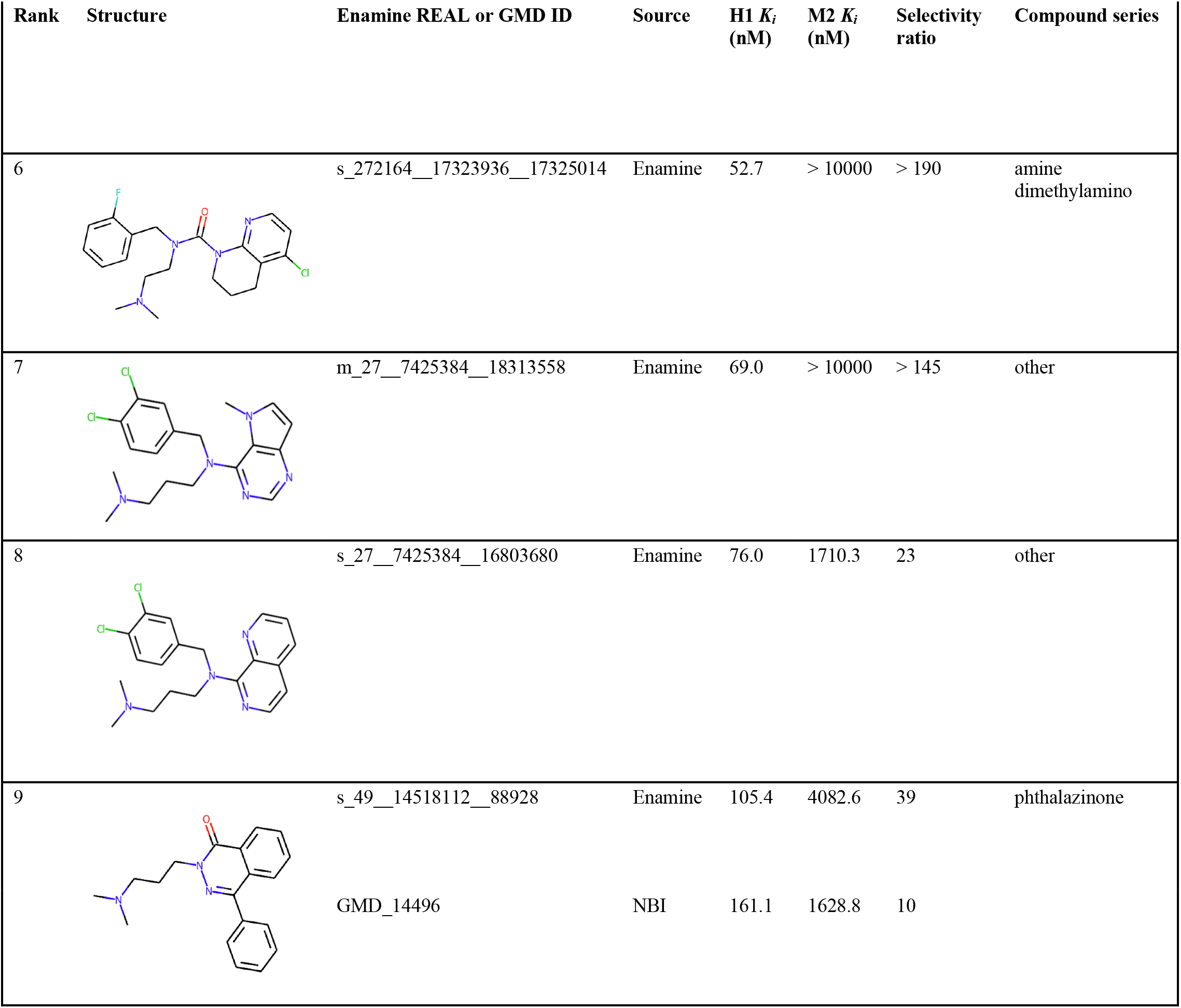
Structures and K_i_ values for most potent H1 binders.

| Rank | Structure | Enamine REAL or GMD ID | Source | H1 $K_i$ (nM) | M2 $K_i$ (nM) | Selectivity ratio | Compound series |
| --- | --- | --- | --- | --- | --- | --- | --- |
| 1 |  | s_49__14518112__1114614 | Enamine | 19.6 | > 10000 | > 509 | phthalazinone |
| 2 |  | s_49__14518112__101877 | Enamine | 19.8 | > 10000 | > 504 | phthalazinone |
|  |  | GMD_12264 | NBI | 27.1 | > 10000 | > 368 |  |
| 3 |  | s_49__14518112__88929 | Enamine | 34.3 | 4378.2 | 128 | phthalazinone |
|  |  | GMD_12503 | NBI | 38.8 | > 10000 | > 258 |  |
| 4 |  | s_27__7100858__10689034 | Enamine | 38.0 | 4839.8 | 127 | benzodiazol |
| 5 |  | GMD_18471 | NBI | 48.0 | 2679.0 | 56 | benzodiazol |

**Table 2. Structures and $K_i$ values for most potent H1 binders (continued)**
| Rank | Structure | Enamine REAL or GMD ID | Source | H1 $K_i$<br>(nM) | M2 $K_i$<br>(nM) | Selectivity<br>ratio | Compound series |
| --- | --- | --- | --- | --- | --- | --- | --- |
| 6 |  | s_272164__17323936__17325014 | Enamine | 52.7 | > 10000 | > 190 | amine<br>dimethylamino |
| 7 |  | m_27__7425384__18313558 | Enamine | 69.0 | > 10000 | > 145 | other |
| 8 |  | s_27__7425384__16803680 | Enamine | 76.0 | 1710.3 | 23 | other |
| 9 |  | s_49__14518112__88928<br>GMD_14496 | Enamine<br>NBI | 105.4<br>161.1 | 4082.6<br>1628.8 | 39<br>10 | phthalazinone |

Of the top nine compounds, four contained a phthalazi-none moiety and three a benzimidazole or benzothiazole group. All of the most potent binders contain a propyl-or ethyldimethylamine chain, which is commonly found in compounds with known anti-H1 activity such as diphenhydramine, chlorpheniramine, and tripelennamine.

### hERG Binding Results

Results from competitive binding assays for the hERG potassium channel are shown for the 9 strongest H1 binders in Table 3 and are plotted in Figure 9 for all 35 compounds with measured hERG binding. The left-hand plot compares the measured *pK_i_* values for binding to H1 and to hERG. The three strongest H1 binders were about 50-to 75-fold selective for H1 over hERG. While some of the compounds tested had hERG *K_i_*’s close to our target of 10 μM, they were all weak H1 binders and were only about 10-to 25-fold selective for H1. The right-hand plot in Figure 9 compares the measured *pIC_50_* values for competitive hERG binding against the *pIC_50_* values predicted by the model used in the GMD optimization. For the majority of compounds, the measured *pIC_50_* was greater than the predicted value. When we examined the model for possible sources of bias, we noted two potentially relevant factors. One was that the *IC_50_* data used for training was actually a mixture of data from binding assays (such as competitive radioligand binding experiments) and functional assays of channel inhibition (such as patch clamp and ion flux assays). The presence of functional data in the training set may have led the model to underpredict the strength of binding to the hERG channel. The other factor is that the compounds used in the hERG training set were structurally dissimilar to the ones used to train the H1 and M2 models, so the hERG model may have been operating far from its domain of applicability. It is possible that the GMD optimization could produce compounds with less hERG binding if a model trained purely on binding data were used for hERG predictions, especially if it incorporated the same hybrid training strategy used for the H1 and M2 models.

**Table 3.** hERG competitive binding results for strongest H1 binders.

| Rank | Enamine REAL or GMD ID | H1 $K_i$ (nM) | hERG $K_i$ (nM) | H1/hERG selectivity |
| --- | --- | --- | --- | --- |
| 1 | s_49__14518112__1114614 | 19.6 | 1486 | 76 |
| 2 | s_49__14518112__101877 | 19.8 | 946 | 48 |
|  | GMD_12264 | 27.1 | 1338 | 49 |
| 3 | s_49__14518112__88929 | 34.3 | 643 | 19 |
|  | GMD_12503 | 38.8 | 866 | 22 |
| 4 | s_27__7100858__10689034 | 38.0 | 1586 | 42 |
| 5 | GMD_18471 | 48.0 | 400 | 8 |
| 6 | s_272164__17323936__17325014 | 52.7 | 300 | 6 |
| 7 | m_27__7425384__18313558 | 69.0 | 604 | 9 |
| 8 | s_27__7425384__16803680 | 76.0 | 65 | 1 |
| 9 | s_49__14518112__88928 | 105.4 | 2621 | 25 |
|  | GMD_14496 | 161.1 | 2529 | 16 |

**Figure 9.**
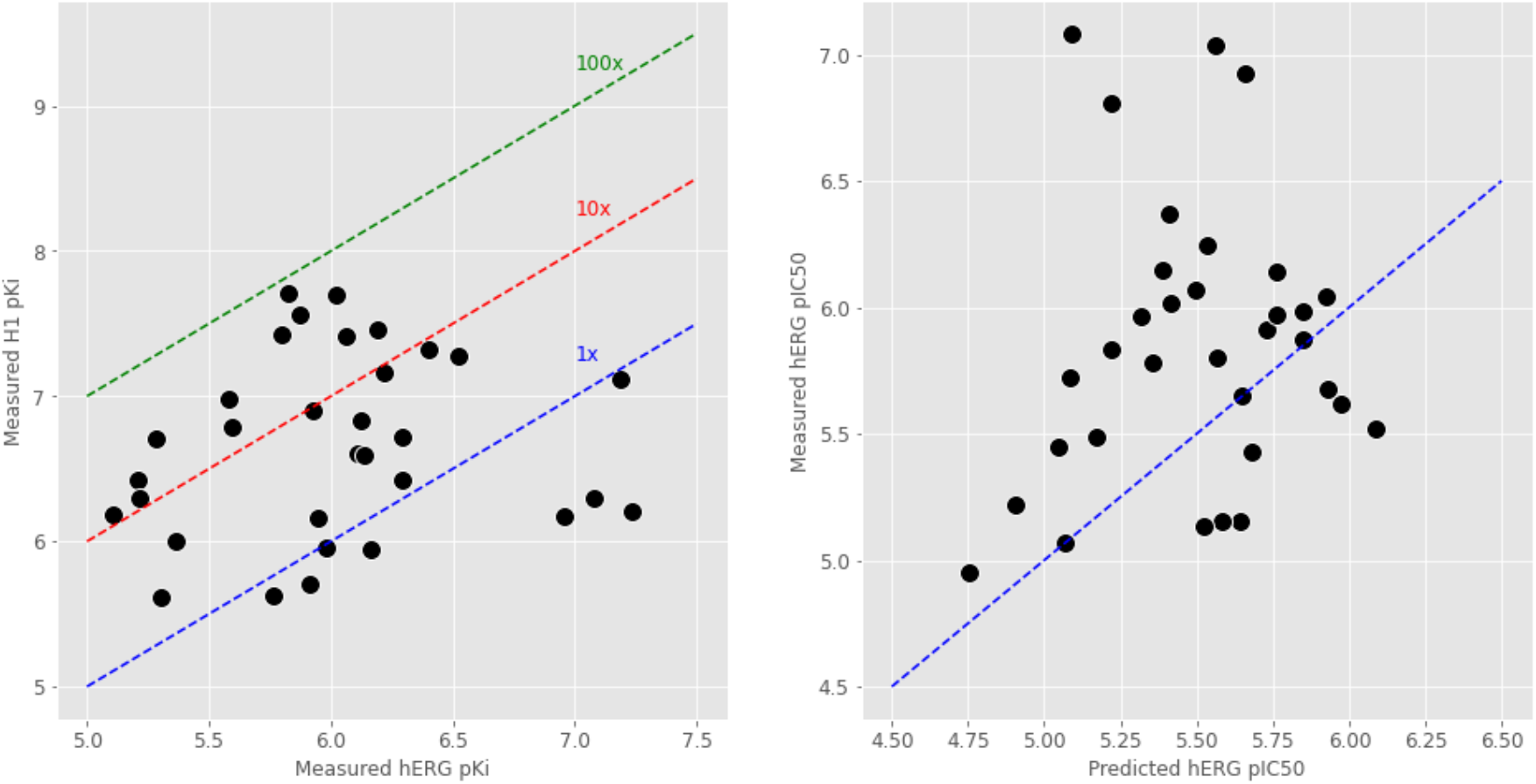
Measured H1 vs hERG binding *pK_i_*; and measured hERG binding *pIC_50_* vs predicted hERG *pIC_50_*

## CONCLUSIONS

In this study we successfully demonstrated the application of the ATOM GMD platform to a realistic drug design problem, namely the optimization of compounds to bind histamine H1 receptors selectively relative to targets associated with common side-effects (M2 and hERG), and satisfying several other properties associated with developability and drug-likeness. Compared to previously published studies of generative molecular design, we synthesized and experimentally tested a somewhat larger number of compounds proposed by our GMD process. Of the 103 distinct molecular structures synthesized, 8 bound H1 with *K_i_*’s between 10 and 100 nM, and 6 were at least 100-fold selective relative to muscarinic M2 receptors.

Several innovations contributed to the success of the project. We found that our use of a genetic algorithm, with JT-VAE latent vectors playing the role of chromosomes, enabled the GMD process to explore a diverse chemical space while focusing on regions associated with desired chemical properties. A flexible and customizable cost function, together with a broad set of property prediction models, allows GMD users to control the importance and stringency of constraints on an unlimited number of design criteria. Including applicability domain indices as cost constraints ensures that the population of proposed molecules remains within the realm where model predictions are reliable, without overly restricting the diversity of compounds. The incorporation of both single- and multiple-concentration activity data into our model training procedure expands the domain of applicability of our models and facilitates an active learning strategy through which compounds can be selected for testing to maximize the expected improvement of model prediction accuracy. Finally, our GMD software platform, based on a scalable, parallelized workflow, allows the evaluation of millions of compounds within 24 hours on a modest-sized compute cluster. When combined with rapid searches of virtual compound databases and scoring of targeted analogs, this platform provides an attractive alternative to brute-force virtual screening.

## EXPERIMENTAL SECTION

### Case Study Definition

To evaluate the performance of the ATOM GMD platform, we applied it to a well-defined lead optimization problem: the design of a potent histamine H1 receptor antagonist with desirable pharmacological properties – in particular, selectivity relative to the muscarinic acetylcholine receptors M1 through M5, and non-inhibition of the human *ether-a-go-go*-related gene (hERG) potassium channel. H1 is well known as a mediator of allergic rhinitis, urticaria and other inflammatory symptoms, as well as arousal in the brain. First generation H1 antihistamines such as diphenhydramine penetrate the blood-brain barrier (BBB), leading to sedation; they also inhibit muscarinic receptors, causing anticholinergic effects such as dry mouth and constipation. Second generation H1 antagonists are more selective with respect to muscarinic receptors and do not pass the BBB; this makes them less sedating, but limits their use when sedation is a desired effect (e.g., when treating insomnia). Two second generation H1 antihistamines, astemizole and terfenadine, were withdrawn from the market due to their inhibition of hERG, resulting in potentially fatal cardiac arrhythmias; consequently, drug candidates are routinely screened for activity against this channel.

For this pilot study, we did not address BBB penetration; we instead focused on optimizing compounds for selective binding to H1, while not binding the muscarinic receptors or acting on hERG. We chose M2 as a representative for the class of muscarinic receptors; this was a reasonable choice because, when we examined publicly available data in which the same compounds were tested against all 5 receptors, binding affinities were strongly correlated between M2 and the other 4 receptors; most M2 binders also bound M1-M5 nonselectively.

### Optimization Criteria

To limit the scale of this case study we proposed an optimization objective function involving three *in vitro* measurable properties and seven calculated attributes. Our specific design criteria are shown in Table 4. The target H1 affinity was chosen to be in the same range as that of existing potent H1 antihistamines; the M2 and hERG targets were selected to yield 1,000-fold and 10,000-fold selectivity, respectively, for compounds meeting the H1 affinity target. Synthetic accessibility scores^21^ are computed using RDKit^22^, and are larger for more complex molecules; the target threshold is in the typical range of catalog-purchasable compounds. The remaining criteria are standard Lipinski rule-of-5 ranges for drug-likeness.

**Table 4.**
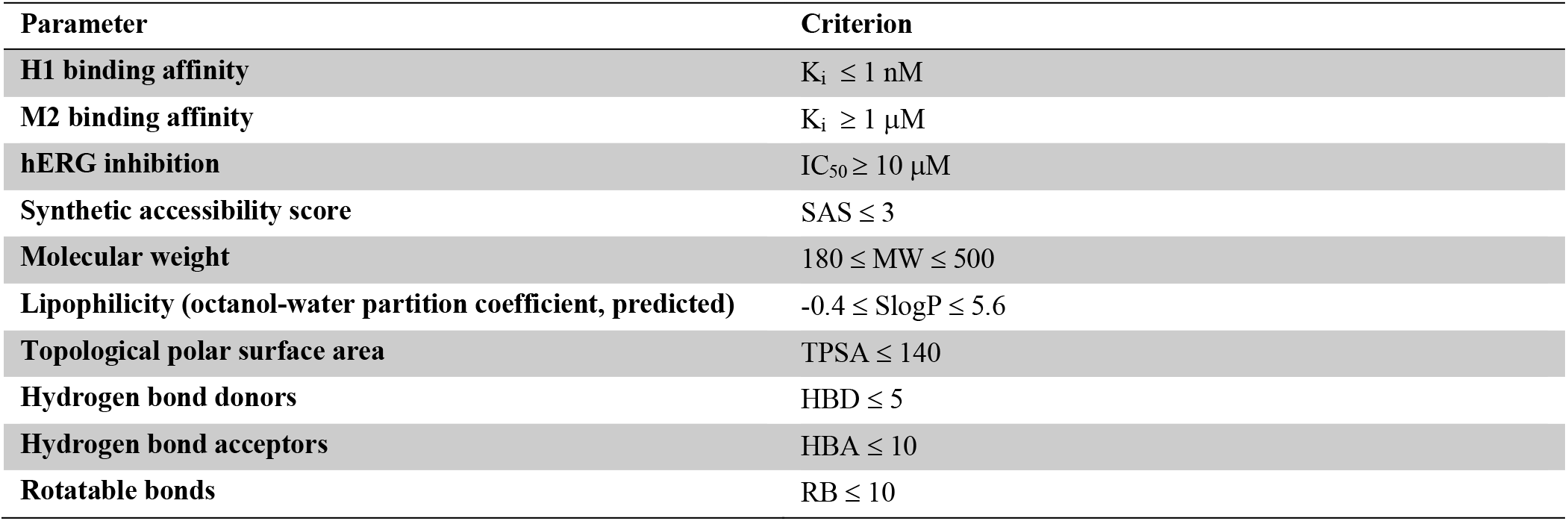
Target design criteria for H1 inhibitor case study.

### Generative Molecular Design Approach

The general approach in the ATOM GMD platform is to propose a population of candidate molecules, predict their functional properties, including efficacy for the specific target, safety liabilities, pharmacokinetics, and developability, then optimize the compound structures in a learned latent representation based on optimization goals for each predicted property. A high-level view of this optimization process is shown in Figure 10. The entire GMD workflow is implemented on a high-performance computing cluster that enables joint optimization of multiple molecular properties for a large population of proposed molecules.

**Figure 10.**
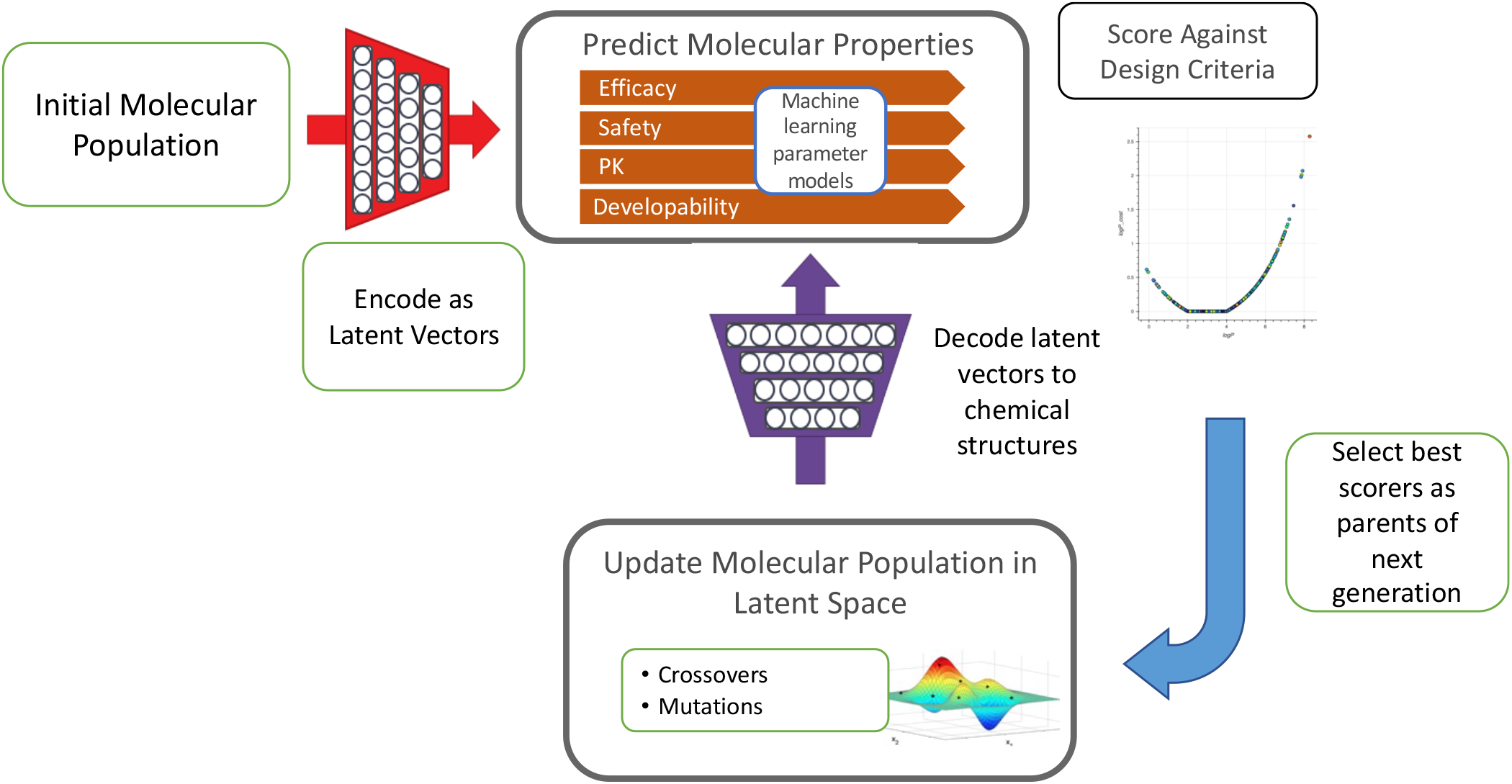
A high-level schematic view of the GMD molecular optimization process. Property predictions for the molecule population are compared to design targets and optimized in a latent space representation.

### Molecular Generative Model

The GMD process incorporates a junction-tree variational autoencoder (JT-VAE) to map a population of molecular structures to a learned latent space. The JT-VAE used here is based on the package developed by Jin, et al^10^; details of the model implementation are given in Supplemental Information. Briefly, the encoder component of the JT-VAE maps each chemical structure into a continuous latent vector consisting of two parts: A 28-dimensional vector encoding a tree of chemical substructures, and a 28-dimensional vector encoding the details of the bonds connecting or shared between substructures. The decoder component of the JT-VAE performs the reverse transformation, from latent vectors to chemical structures. During training, the JT-VAE learns a “vocabulary” of chemical substructures present among the training compounds; generated molecules are limited to those that can be constructed from the vocabulary. Therefore, a training set can be selected to bias the generated structures toward families of compounds with known activity against a desired target; or can be selected for greater diversity to allow the generative model to explore a broader chemical space.

In this project a combination of the two training approaches was used to balance diversity and specificity. An initial set of 1825 compounds provided by Neurocrine Biosciences, restricted to those with >90% receptor binding at 10 uM, was combined with molecules selected from the ChEMBL 28^23^ and GoStar^24^ databases having measured H1 *pK_i_* values greater than 7. These molecules were provided as queries to the Rapid Isostere Discovery Engine (RIDE)^25^, part of the ICM-Pro software^26^. The RIDE engine searched the Enamine REAL database^27^ for molecules with similar shapes and matching atomic property fields^28,29^ to the query molecules. The resulting set of 24,741 molecules was split into a 23,741 compound training set and a 1,000 compound evaluation set. A hyperparameter search training JT-VAE models with 110 different parameter configurations was performed using the JT-VAE jtnn_hyperparam tool; the best performing model, as determined by reconstruction accuracy on the evaluation set, was used for all subsequent GMD runs. Parameters for this model are shown in Supplemental Table 3.

### Property Prediction Models

Quantitative structure activity relationship (QSAR) models were trained to predict H1 and M2 binding affinities and hERG inhibition using the ATOM Modeling PipeLine (AMPL) ^0^. The training data for the H1 and M2 models were obtained by combining *pK_i_* binding data from the ChEMBL 28 and GoStar databases with single-concentration percent binding measurements provided by Neurocrine. The hERG model was trained with *pIC_50_* data from the ChEMBL, Excape^31^ and Drug Target Commons^32^ databases. For all data sets, we used the RDKit and MolVS^33^ Python packages to standardize SMILES strings and remove salts. The data for the H1, M2 and hERG models were partitioned into training, validation and test sets using the “scaffold” splitter implemented in the DeepChem package^34^. Full details of the data sources, compound counts and split proportions are given in Supplemental Tables 1 and 2.

SMILES strings from the combined dataset were mapped to chemical descriptors using the Molecular Operating Environment (MOE) software^35^. The complete set of MOE descriptors was pruned to exclude those that were undefined or uncomputable for most compounds in our training set, yielding 306 computable descriptors for each compound. Spurious correlations between pairs of descriptors that increase with molecule size were eliminated by dividing those descriptors by total atom counts. The resulting descriptors were used as input features for predictive models of H1 and M2 receptor binding and hERG inhibition. Six descriptors were used directly in the final cost function, as shown in Table 4.

To predict H1 and M2 binding and hERG inhibition, fully connected neural networks with between one and three hidden layers were trained using hyperparameter searches to select the optimal network architecture, learning rate and dropout parameters for each target and feature set. The models were trained using an early stopping algorithm, in which training was terminated when the coefficient of determination *R^2^* over the validation set failed to improve after 100 training epochs; the model parameters yielding the maximum *R^2^* value were then saved for later predictions. The parameters for the final selected models are shown in Supplemental Tables 1 and 2.

Training the H1 and M2 binding models presented a special problem because the training data contained a mixture of *pK_i_* values and single-concentration percent binding values from competitive radioligand binding experiments. It was important to include the single concentration measurements because they covered a unique region of chemical space not represented in the *pK_i_* dataset; but these measurements tended to be noisy, with the noise increasing for weaker binders (for which the observed radioligand binding is greater). We addressed this issue by developing a hybrid training procedure, in which the model predicts *pK_i_* values first, then infers percent binding at specific concentrations using the Cheng-Prusoff relation. The loss function minimized during training combines different types of terms for the two types of training data points: a standard *L_2_* loss for the *pK_i_* values and a Poisson loss term for the percent binding values. Details about the hybrid loss function and training procedure are provided in Supplemental Information. The hybrid model was implemented with PyTorch and has been incorporated into the AMPL toolkit.

During initial runs of the GMD pipeline, we found that applicability domain (AD) constraints were necessary to prevent the optimization process from favoring proposed compounds that were chemically too dissimilar from the compounds in the H1 and M2 model training datasets, so that the model predictions were unreliable. Following a method described by Mathea et al^36^, we implemented a Z-score AD index for each model and calibrated it using logistic regression to yield an estimated probability for each model to generate predictions outside the range of its training data, given an input chemical structure. Details of the AD index algorithm are given in Supplemental Information

### Objective Function Implementation

The GMD optimization process begins with an initial working population of molecules and iteratively updates it over several hundred generations, with the goal of evolving a set of compounds that minimize an objective cost function. The cost function is defined explicitly in terms of molecular property target values, allowing a drug discovery scientist to trade off different properties in order to maximize a therapeutic window, e.g. finding compounds for which the minimum effective concentration is well below the maximum safe concentration in the target patient population. We used a simple yet fully customizable scoring function to assign costs to the predicted properties. The cost function for a molecule is calculated as a weighted sum of exponential functions of the distances between the target value and the predicted value for each property evaluated; the weights, scale factors, functional forms and target ranges can be individually specified for each property. A detailed list of the cost function terms for our selective H1 inhibitor GMD runs is given in Supplemental Information.

### Genetic Algorithm Optimizer

We implemented a genetic algorithm-based optimizer to search the latent space for optimal molecules, treating latent vectors analogously to chromosomes and vector elements to genes, and using crossovers and mutations to generate new molecules. Candidate parents for crossover were chosen by tournament selection, randomly sampling sets of three latent vectors and selecting the top scorer from each set. Elements of parent vector pairs were then sampled randomly to produce a number of offspring vectors equivalent to 50% of the original population. We note that the population of latent vectors in any generation may include several that decode to the same SMILES string. To increase diversity, we limited to three the number of latent vectors per generation encoding the same SMILES string that could be selected for crossover.

Subsequently the optimizer applied a combination of global random mutations and single-point fixed-step local mutations on the top candidates in each generation. The global mutation step involved randomly selecting 10% of the working population of latent vectors, choosing 20% of the elements at random in each vector, and adding to each selected element a random perturbation value sampled from a normal distribution. The fixed-step mutation was performed by selecting the best scoring 10% of latent vectors and adding a perturbation of fixed magnitude (0.1) and random sign to a randomly selected element of each vector. Although most variational autoencoders are optimized to generate latent vectors with standard normal distributions for each element, the two classes of elements (encoding the junction tree and molecular graph) of latent vectors produced by the JT-VAE follow normal distributions with different standard deviations. Therefore, we multiplied both the random and fixed-step perturbations by the appropriate standard deviation, depending on which type of element was being mutated.

The new latent vectors generated by crossover and mutation were then combined with the remaining unselected vectors to produce the source population for the next generation. The loop was terminated when it reached a specified number of generations or after a specified execution time.

### High Performance Compute Framework

The ATOM GMD process is parallelized for high-performance computing clusters and scales both to arbitrary candidate molecule population sizes and to any number of property prediction models applied in the optimization loop. To facilitate the computationally intensive operations needed to generate, evaluate, and optimize a chemical library against the desired properties, we built a flexible, scalable pipeline framework, called SPL, using a RabbitMQ AMQP message broker^37^ to coordinate a configurable set of distributed, object-oriented workers. The workers perform the featurization, prediction, encoding/decoding, optimization, and batching/aggregation tasks required to perform generative molecular design at large scale, as diagrammed in Figure 11. The pipeline can be configured with different workers to provide each of these functions, allowing the researcher to experiment with different generative models, featurizers, optimizers and so on. The SPL framework facilitates communication and scaling across heterogeneous compute nodes, providing efficient batching, distribution, parallelization, and supervision of work to utilize the available computational resources. It supports scheduling and resource management using workload managers such as Slurm^38^ and Flux^39^. The instance of the framework used for the case study reported here utilized 34 nodes (with 48 CPU cores per node) on the Corona supercomputer at Lawrence Livermore National Laboratory; other ATOM GMD design projects have been distributed across hundreds of nodes spanning multiple clusters.

**Figure 11.**
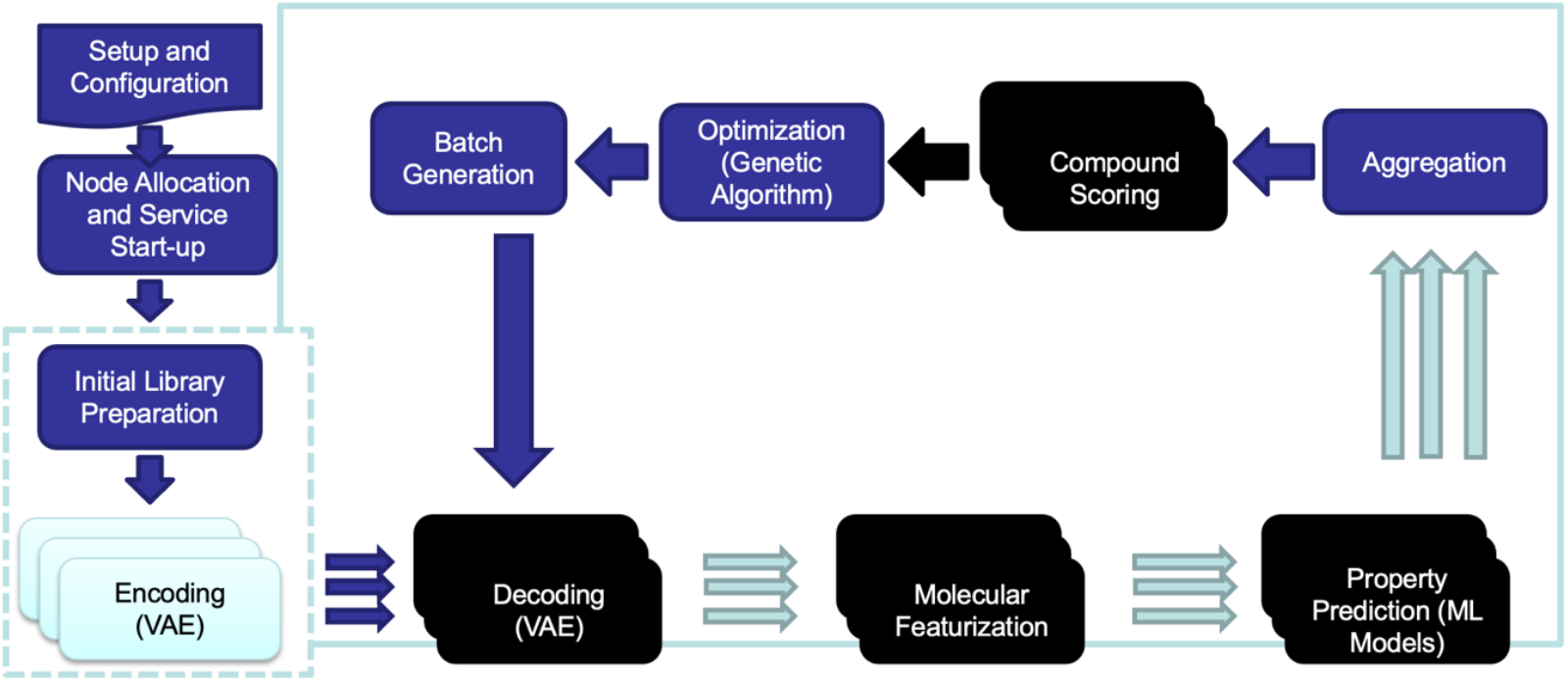
High performance compute workflow overview for the ATOM Scalable Pipeline Loop (SPL)

### GMD Runs and Parameters

Two GMD optimization runs were performed to generate compounds for experimental validation, a small-scale initial run and a large-scale production run. The main differences between the two runs were the sizes of the initial compound sets, the numbers of generations and the models used for H1 and M2 *pK_i_* predictions. Both initial compound sets were selected from the sets of molecules used to train and test the junction tree variational autoencoder. The first GMD run was initialized with the 1,000 compound evaluation set described earlier and ran for 200 generations; the second run was initialized with the full set of 24,741 compounds and ran for 253 generations. For the second run, the H1 and M2 models were replaced with updated versions trained on datasets incorporating new experimental data provided by Neurocrine, consisting of competitive binding measurements for 160 compounds at 10 nM concentration.

### Compound Selection for Experimental Validation

The output from a GMD run is a table of compounds proposed and evaluated in each generation, listing the SMILES string, cost score, individual model predictions and cost score components for each molecule. For each GMD run, we filtered the output table to include only the first appearance of each SMILES string, then selected the 2,000 best scoring molecules as candidates for validation. For the second run, we clustered the top scoring molecules using Butina’s algorithm with a threshold Tanimoto distance of 0.5, then selected the best scoring representative from each cluster, producing a set of 566 candidate molecules. The two candidate sets were combined to produce a pool of 2,566 candidates for synthesis and experimental validation.

To obtain additional compounds that could be purchased rather than requiring custom synthesis in our laboratory, and to improve our predictive models by testing a more structurally diverse set of compounds, we searched for near matches to the top-scoring 2,000 molecules from each run in the 13.5 billion compound Enamine REAL virtual library^27^. The search was performed using the Arthor similarity search software version 3.4^40^ For each GMD compound, we selected the closest matching (smallest Tanimoto distance) compound from Enamine REAL. We then scored the Enamine compounds using the GMD pipeline, and selected the Enamine matches that scored at least well as the 2,000^th^ best scoring compound from the original GMD run. This resulted in 38 high scoring matches from the first GMD run (including 3 exact matches) and 57 from the second, yielding 95 total compounds that were ordered from Enamine.

We then selected a group of compounds for custom synthesis from the original 2,566 compound pool, based on input from Neurocrine chemists. We prioritized three groups of GMD compounds that shared common scaffolds and therefore could be synthesized efficiently, by leveraging common precursors and intermediate steps, and potentially making use of Neurocrine’s automated synthesis platform. These three chemical series and associated SMARTS patterns and structures are shown in Table 5.

**Table 5.**
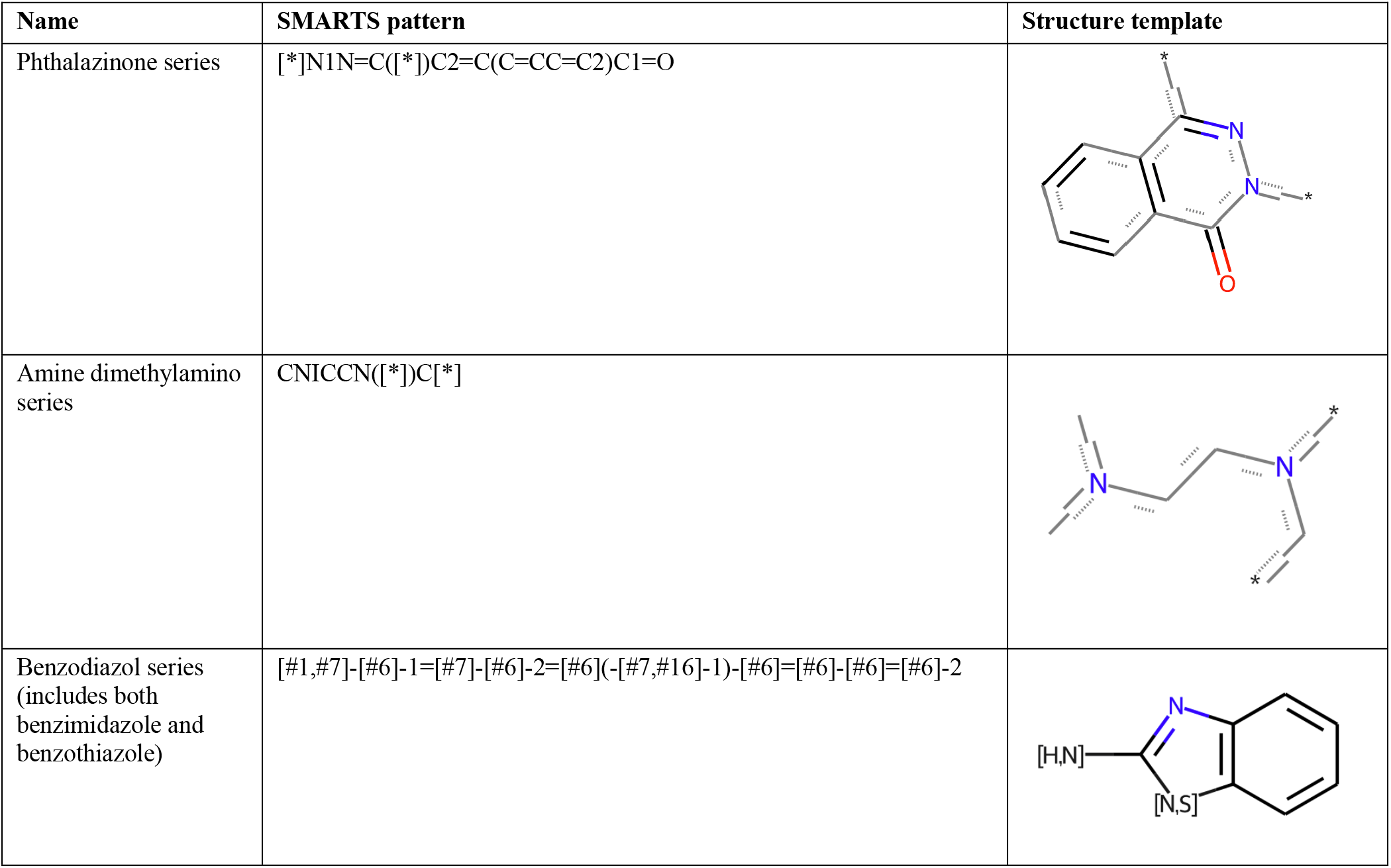
Chemical series selected for synthesis at Neurocrine.

| Name | SMARTS pattern | Structure template |
| --- | --- | --- |
| Phthalazinone series | [*]N1N=C([*])C2=C(C=CC=C2)C1=O |  |
| Amine dimethylamino series | CN(C)CCN([*])C[*] |  |
| Benzodiazol series (includes both benzimidazole and benzothiazole) | [#1,#7]-[#6]-1=[#7]-[#6]-2=[#6](-[#7,#16]-1)-[#6]=[#6]-[#6]=[#6]-2 |  |

We eliminated molecules from series already known to inhibit H1, such as tricyclics; with reactive or unstable moieties; with multiple stereoisomers; and with structures that were difficult to synthesize, such as quaternary centers, tetrasubstituted olefins or ambiguous olefin geometry. The Molecule.One retrosynthesis software^41^ was used to develop synthesis plans and identify molecules from the candidate pool that could be created with the fewest steps. In all, 48 compounds proposed by the GMD runs were selected for custom synthesis. Seven additional compounds belonging to the phthalazinone series were designed manually and added to the synthesis set, resulting in a total of 55 compounds planned for synthesis at Neurocrine and 95 to be purchased from Enamine. The three compounds that had exact matches in the Enamine REAL catalog, all of which shared the phthalazinone scaffold, were synthesized both at Neurocrine and by Enamine, allowing us to check the consistency of results between the three pairs of synthesis products.

### Competitive Radioligand Binding Assays

All synthesized compounds were tested for binding to histamine (H1), muscarinic (M2) and hERG receptors through radioligand competitive binding assays at Neurocrine Biosciences, Inc. All compounds were assayed in 96-well plates in single point with final concentrations at 10μM. Positive controls were serially diluted on each plate for data validation and normalization. 50 μL radioligand, 25 μL compound and 75 μL membrane with overexpressed receptor were added to 96-well white round bottom polystyrene NBS microplates (Corning #3605) and incubated on 25°C plate warmer for 90 minutes. Plates were then filtered on GF/B filter plates precoated with 0.1% PEI (PerkinElmer #6005177) and washed with 1.2mL wash buffer and fan dried. Scintillation fluid was added and plates were incubated overnight. Finally, plates were read on a PerkinElmer TopCount reader.

The radioligand competition binding data were analyzed at Neurocrine Biosciences, Inc. using the R programming language^42^. For each 96-well plate, the positive control data were fit to a 4-parameter logistic curve. Resulting top and bottom asymptotes from the regression fit were used to calculate a normalized percentage of receptors bound to compound for each well using the following equation:

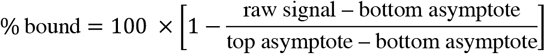

The normalized percent binding data with compound structures were subsequently delivered to ATOM for machine learning model development.

Eight-point dilution series were performed for compounds with greater than 50% binding at 10 μM; inhibition constants *K_i_* and *IC_50_*’s were calculated from 4-parameter logistic models in R.

## Notes

Disclaimer: This document was prepared as an account of work sponsored by an agency of the United States government. Neither the United States government nor Lawrence Livermore National Security, LLC, nor any of their employees makes any warranty, expressed or implied, or assumes any legal liability or responsibility for the accuracy, completeness, or usefulness of any information, apparatus, product, or process disclosed, or represents that its use would not infringe privately owned rights. Reference herein to any specific commercial product, process, or service by trade name, trademark, manufacturer, or otherwise does not necessarily constitute or imply its endorsement, recommendation, or favoring by the United States government or Lawrence Livermore National Security, LLC. The views and opinions of authors expressed herein do not necessarily state or reflect those of the United States government or Lawrence Livermore National Security, LLC, and shall not be used for advertising or product endorsement purposes.

## ABBREVIATIONS

AD: applicability domain
AMPL: ATOM Modeling PipeLine
ATOM: Accelerating Therapeutics for Opportunities in Medicine
BBB: blood-brain barrier
ECFP: extended connectivity fingerprint
GAN: generative adversarial network
GMD: generative molecular design
hERG: human ether-a-go-go-related gene
JT-VAE: junction tree variational autoencoder
MOE: Molecular Operating Environment
NBI: Neurocrine Biosciences Inc.
QSAR: quantitative structure activity relationship
RIDE: Rapid Isostere Discovery Engine
t-SNE: t-distributed stochastic network embedding
VAE: variational autoencoder

## SUPPLEMENTARY METHODS AND RESULTS

### Predictive model training data sets

Data for training predictive models of binding affinity for H1 and M2 were obtained from multiple sources:

- Single-concentration competitive radioligand binding measurements contributed by Neurocrine Biosciences, Inc. to the NCI Model & Data Clearinghouse (MODAC); available at https://modac.cancer.gov/searchTab?dme_data_id=NCI-DME-MS01-6611544 and https://modac.cancer.gov/searchTab?dme_data_id=NCI-DME-MS01-7425272.
- *pK_i_* data from the public ChEMBL 28 database and the proprietary GoStar database (Excelra, 2021).

Details on the training data sets are given in Supplementary Table S1.

Data for training the predictive model for hERG inhibition was obtained from ChEMBL 27 and the publicly available Drug Target Commons (Tang et al., 2018) and ExCAPE-DB (Sun et al., 2017) databases. Details of the training set composition are given in Supplementary Table S2.

**Supplementary Table S1.** Data for training predictive models for binding affinities derived from Neurocrine contributed measurements, ChEMBL 28 and GoStar; training/validation/test set splits; and model parameters.

| Property | H1 $pK_i$ dataset | M2 $pK_i$ dataset |
| --- | --- | --- |
| Neurocrine %binding @ 10 μM | 1825 | 1824 |
| Neurocrine %binding @ 10 nM | 160 | 160 |
| ChEMBL $pK_i$ unique compounds | 1214 | 1145 |
| GoStar $pK_i$ unique compounds | 1873 | 655 |
| Overlap between Neurocrine and ChEMBL/GoStar compound sets | 100 | 28 |
| **Total unique compounds** | 4812* | 3596* |
| train/validation/ test split | 80/10/10 | 80/10/10 |
| **Best model parameters:** |  |  |
| Layer architecture | 256 x 16 | 128 x 64 |
| Dropout fraction | 0.3 | 0.3 |
| Learning rate | 5e-05 | 7e-04 |
| Training epochs | 11 | 50 |
| **Performance metrics:** |  |  |
| Coefficient of variation ($R^2$) | 0.52 | 0.75 |
| Pearson correlation coefficient | 0.74 | 0.87 |

\*Total compound count is less than sum of components due to overlap between Neurocrine and ChEMBL/GoStar compound sets.

**Supplementary Table S2.** Data for training predictive model for hERG inhibition derived from ChEMBL 27, Drug Target Commons and Excape databases; training/validation/test set splits; and model parameters.

| Property | Value |
| --- | --- |
| ChEMBL 27 $pIC_{50}$ unique compounds | 6394 |
| Drug Target Commons $pIC_{50}$ unique compounds | 2301 |
| ExCAPE-DB $pIC_{50}$ unique compounds | 3225 |
| **Total unique compounds** | **11,920** |
| train/validation/test split percentages (for hyperparameter search) | 70/15/15 |
| train/validation split (for final model, using best hyperparameters) | 85/15 |
| **Best model parameters:** |  |
| Layer architecture | 256 x 64 x 32 |
| Dropout fraction | 0.3 |
| Learning rate | 7e-04 |
| Training epochs | 113 |

### Generative Network Training for the H1 Inhibitor Design

The generative network used for this study was a junction-tree based variational autoencoder (JT-VAE) based on the package developed by Jin, et al (Jin et al., 2018). Our implementation of the JT-VAE was essentially the same as the “fast_jtnn” variant released on the author’s GitHub site (Jin, 2018/2021), except that we ported the code from Python 2.7 to Python 3.6. To establish the latent space representation for the H1 inhibitor chemical design space, we trained a JT-VAE on a set of 24,741 compounds derived by querying the Enamine REAL virtual library for molecules that were isosteric to a seed set of known H1 inhibitors, using the Rapid Isostere Discovery Engine tool in the Molsoft ICM-Pro software. We split the resulting compound library randomly into a 23,741 compound training set and a 1,000 compound test set.

Prior to training, SMILES representations of library compounds were salt-stripped and standardized using the MolVS and RDKit Python packages. We derived the vocabulary set and preprocessed the training library according to the procedures documented on the JT-VAE GitHub site. We trained the JT-VAE using the vae_train.py script with the parameters shown in Supplementary Table 3. Checkpoint files were saved after every epoch. The checkpoint files allowed us to evaluate the performance of the JT-VAE model over the course of training, by loading the model from each checkpoint and using it to encode and decode structures from the held-out test compound set. We evaluated performance using two metrics: The recovery rate (i.e., the fraction of encoded input structures that were recovered perfectly when decoded), and the average Tanimoto similarity between the encoded and decoded structures. For our training run, both metrics reached maximum values at 23 epochs. At the peak, the recovery rate was 7.7% and the mean Tanimoto similarity was 0.524. We used the trained JT-VAE network from this best performing checkpoint as our generative model for all of our subsequent experiments.

**Supplementary Table S3.**
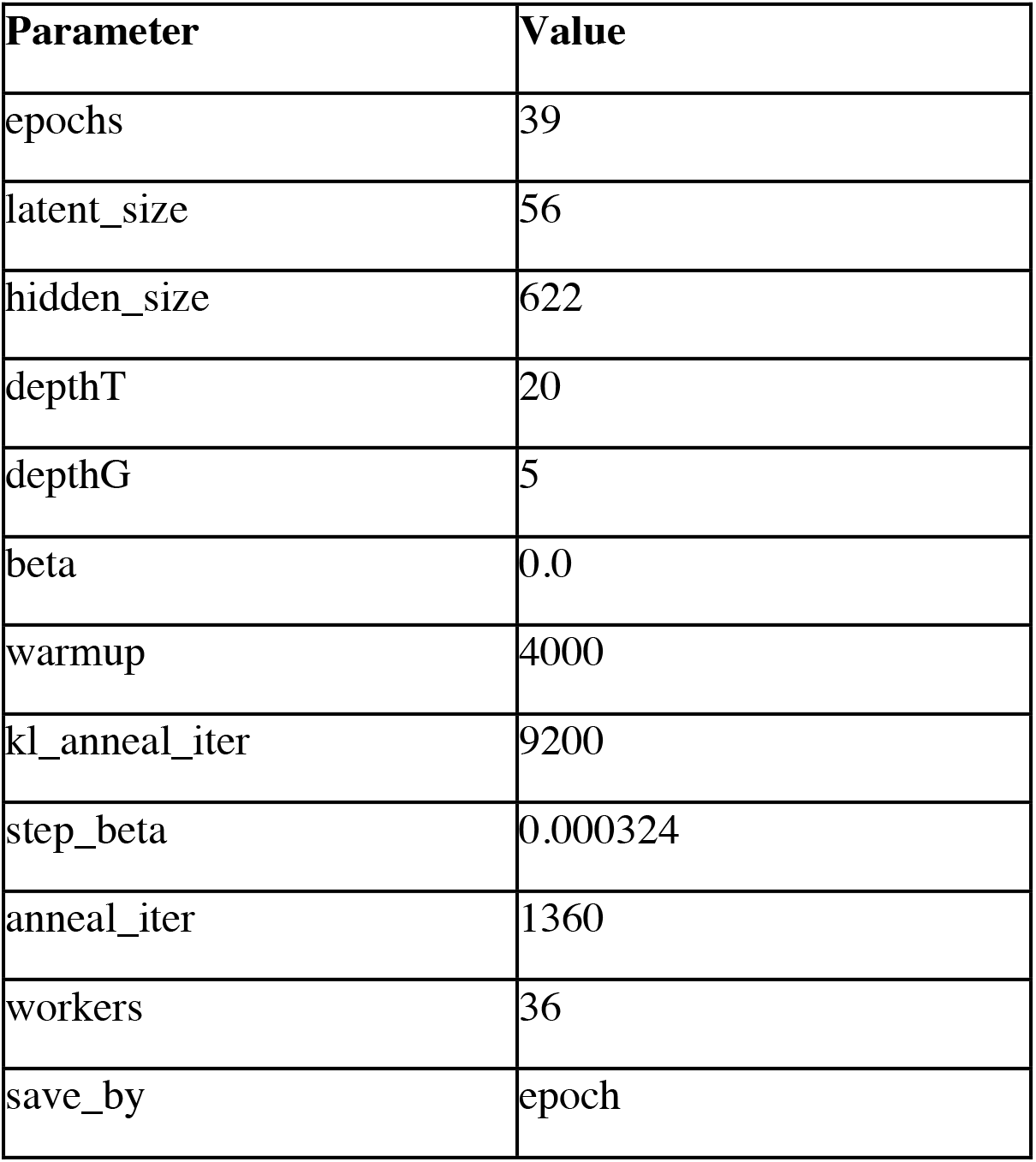
vae_train.py parameters used for JT-VAE training

| Parameter | Value |
| --- | --- |
| epochs | 39 |
| latent_size | 56 |
| hidden_size | 622 |
| depthT | 20 |
| depthG | 5 |
| beta | 0.0 |
| warmup | 4000 |
| kl_anneal_iter | 9200 |
| step_beta | 0.000324 |
| anneal_iter | 1360 |
| workers | 36 |
| save_by | epoch |

### Hybrid model training procedure for H1 and M2 binding prediction

The training data for our H1 and M2 receptor binding models contained *pK_i_* values obtained from public and proprietary data sources along with single-concentration bound fraction values from competitive radioligand binding experiments performed at Neurocrine. Typically, data-driven models to predict receptor binding are trained on *pK_i_* or *pIC_50_* data only, as these values are estimated by fitting logistic curves to multiple percent binding measurements and therefore tend to be less noisy. However, for our project we wished to incorporate the single-concentration measurements into model training, because the compounds measured belonged to a unique set of chemical families not well represented in the *pK_i_* dataset, and we wanted our models to perform well on similar compounds. To make this possible, we developed a hybrid neural network model training procedure that leverages both types of binding data.

To train a hybrid model, we prepare a data table containing columns of compound identifiers, SMILES strings, activities and concentrations. The activity column contains both bound fraction and *pK_i_* measurements; corresponding concentration values are provided for the bound fraction values, while for the *pK_i_* values the concentration field is left blank. The data for both types of measurements are shuffled together randomly, and the dataset is divided into training, validation and test subsets using standard scaffold splits.

The SMILES strings for the compounds are used to compute MOE descriptors as input features for the model.

During training, the feature vectors are input to a multilayer perceptron neural network with weights and biases initialized to Gaussian random numbers. The output from the neural network is treated as a prediction of the *pK_i_* of the input compound. The prediction is then used to compute a term of a loss function, in a way that depends on the type of the corresponding activity data. If the activity is a *pK_i_* value, a standard *L*_2_ loss is computed, equal to the square of the difference between the predicted and measured *pK_i_*. If the activity is a percent binding at some concentration, the model uses the predicted *pK_i_* together with the Cheng-Prusoff relation to estimate the IC_50_, and then the expected fraction of receptors bound to radioligand (rather than the tested compound) at that concentration:

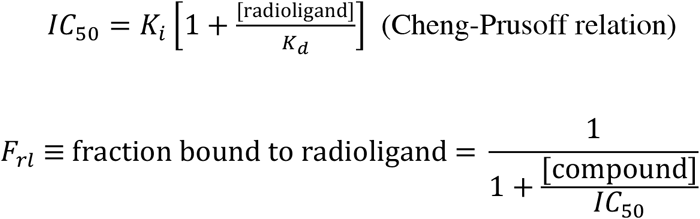

where the *K_d_* in the Cheng-Prusoff relation is the dissociation constant for the radioligand used in the competitive binding experiment. The predicted *F_rl_* value is then compared to the measured *F_rl_* (i.e., one minus the fraction of receptors bound to the test compound) to compute a loss term. Because the measurements in competitive radioligand binding experiments are based on scintillation counting of the residual radioactivity, the noise in these measurements is approximately Poisson distributed; notably, the noise variance scales with *F_rl_*, and therefore increases the more weakly the test compound binds to the receptor. Therefore, the appropriate loss term is a Poisson loss:

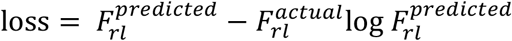

The two sets of loss terms are summed over the measurements in each training batch to create a loss function. The weights and biases in the neural network are then adjusted during training with the goal of minimizing the loss function, using a standard backpropagation algorithm.

### Applicability domain index calculation and calibration

The GMD optimization cost function includes terms that penalize proposed molecular structures that are overly dissimilar to any compounds in the H1 and M2 model training sets, using an applicability domain index (ADI) based on a method described by Mathea et al. (Mathea et al., 2016). The ADI computation can be summarized as follows:

- Compute feature vectors *x_i_* for all compounds in the training set, shifted and scaled so that each feature has mean 0 and standard deviation 1.
- Compute Euclidean distances *d_ij_* = *d*(*x_i_*, *x_j_*) between all pairs of feature vectors.
- For each compound *I* in the training set, calculate the mean distance to its *K* nearest neighbors in the training set. *K* was set to 5 for our models.
- Compute the mean 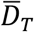 and standard deviation *σ_D_* of the mean distances.
- For each proposed compound *p*, compute a scaled and shifted feature vector and calculate the mean distance 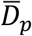 to its *K* nearest neighbors in the training set.
- Calculate a *Z*-score for compound 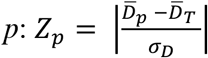.

The *Z*-score by itself can be used to rank proposed compounds according to their dissimilarity to an individual model’s training set. However, the range of *Z*-score values associated with reliable *pK_i_* predictions was found to vary greatly between the H1 and M2 models. To use the *Z*-score to assign a cost penalty to overly dissimilar compounds required a further calibration step. We noted that, if we binned the compounds proposed by an initial GMD run by *Z*-score and plotted the fraction of compounds with predicted *pK_i_* outside the range of the training data (greater than 10.3 for the H1 model and less than 3.7 for the M2 model), the fractions roughly followed a logistic curve, as shown in Figure S1. Accordingly, we fit logistic regression models to the model predictions for H1 and M2, with Z-score as the predictor variable and an indicator variable (H1 *pK_i_* > 10.3 or M2 *pK_i_* < 3.7) as the response. The calibrated AD index is the probability computed by the fitted logistic regression model.

**Figure S1:**
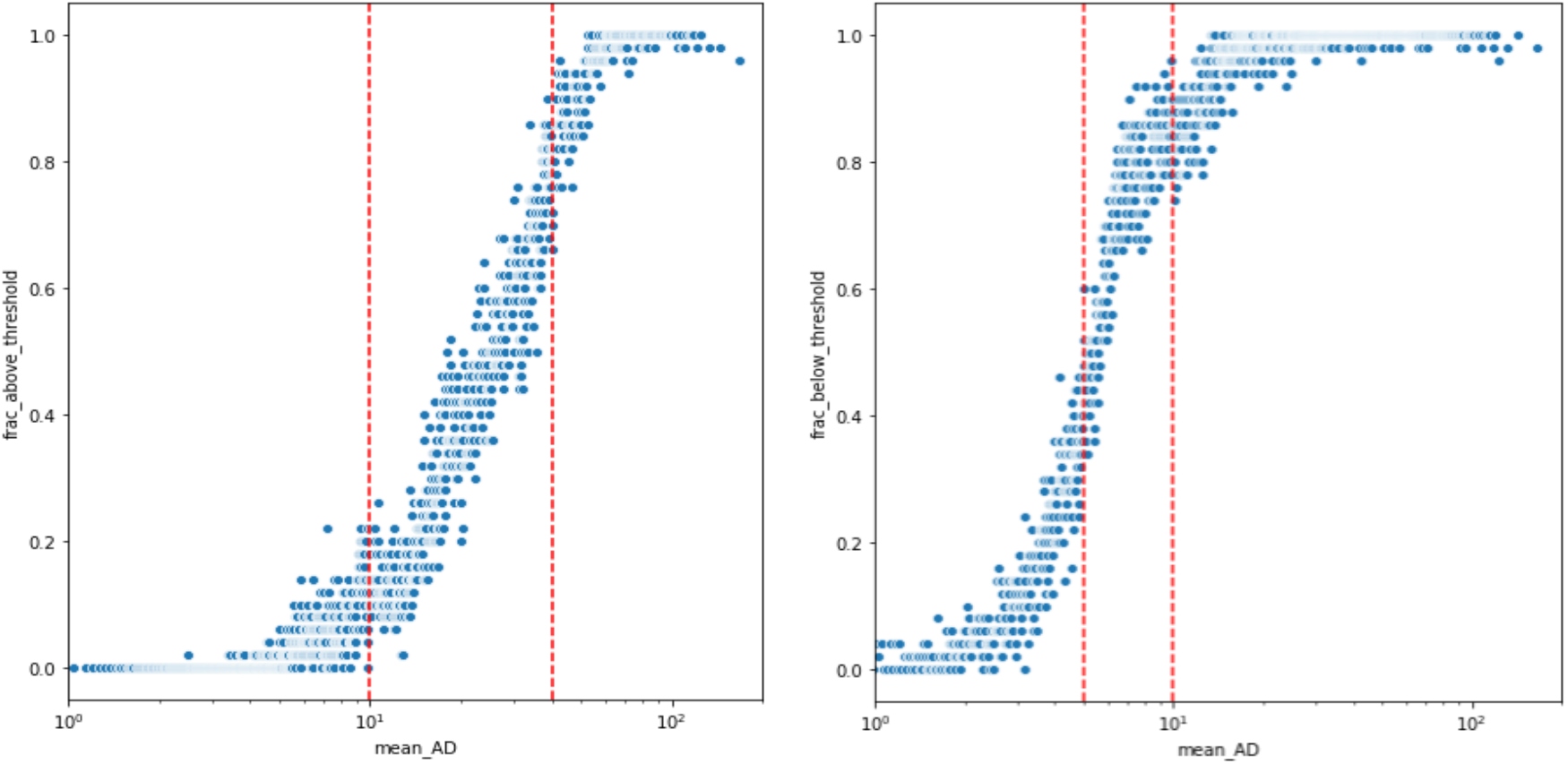
Fraction of compounds proposed in initial test GMD run with predicted H1 (left) and M2 (right) pK_i_ values outside the training data range in bins of 50 compounds grouped by Z-score AD index.

### Cost function for selective H1 inhibitor optimization

The cost function used to score molecules proposed by the GMD software is a weighted sum of penalty terms based on 12 predicted properties:

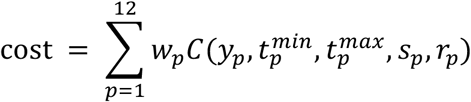

where *y_p_* is the predicted value for property *p*; 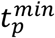 and 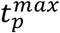 are the target minimum and maximum values for property *p*; ***s_p_*** is a property-specific scale factor; ***r_p_*** is a binary value indicating whether the term value must be rectified (forced to be non-negative); and ***w_p_*** is a weight factor. Either 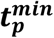 or 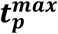 may be left unspecified, in which case the values are treated as −∞ and +∞ respectively. The penalty term has the general form:

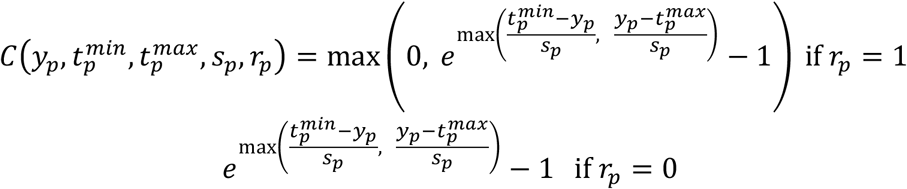

**Supplementary Table S4.**
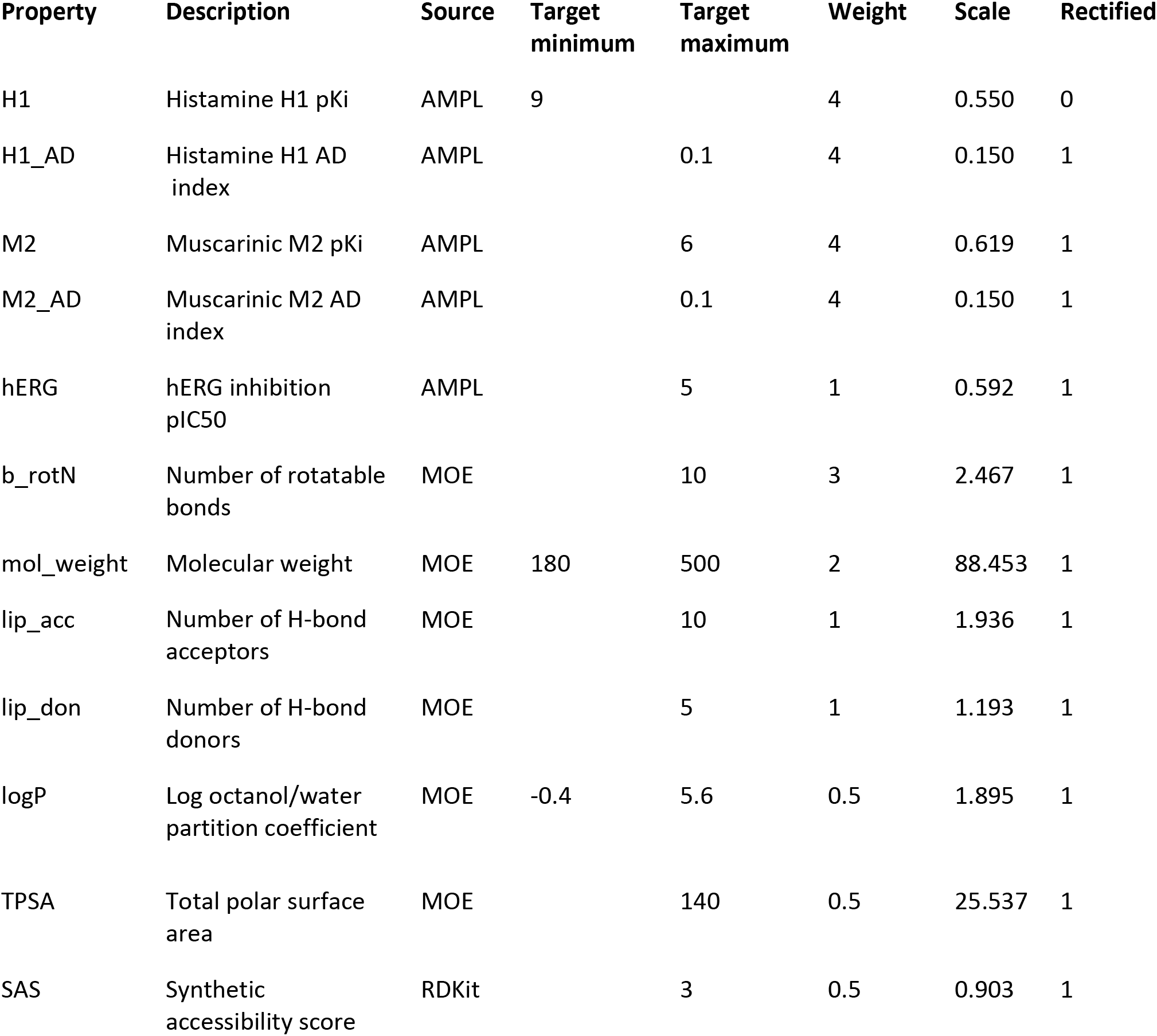
Properties to be optimized and associated cost function parameters

The rectified penalty term for log *P*, in which the target range is bounded on both sides, is plotted in Figure S2. In general, the penalty is zero for property values within the target range, and increases steeply with the distance of the predicted value from the adjacent end of the range. The scale factor *s_p_* determines the steepness of the exponential curve. By default *s_p_* is set to the standard deviation of property *p*’s values in the predictive model training set; GMD users can adjust this value, along with the weight *w_p_*, to assign steeper penalties for properties that need to be more tightly controlled. Generally most penalty terms are rectified, but rectification can be turned off for properties such as on-target potency; this allows the GMD process to permit highly potent compounds with a few liabilities to go forward for further optimization. The cost function parameters used for the H1 inhibitor study are listed in Supplementary Table S4.

**Figure S2:**
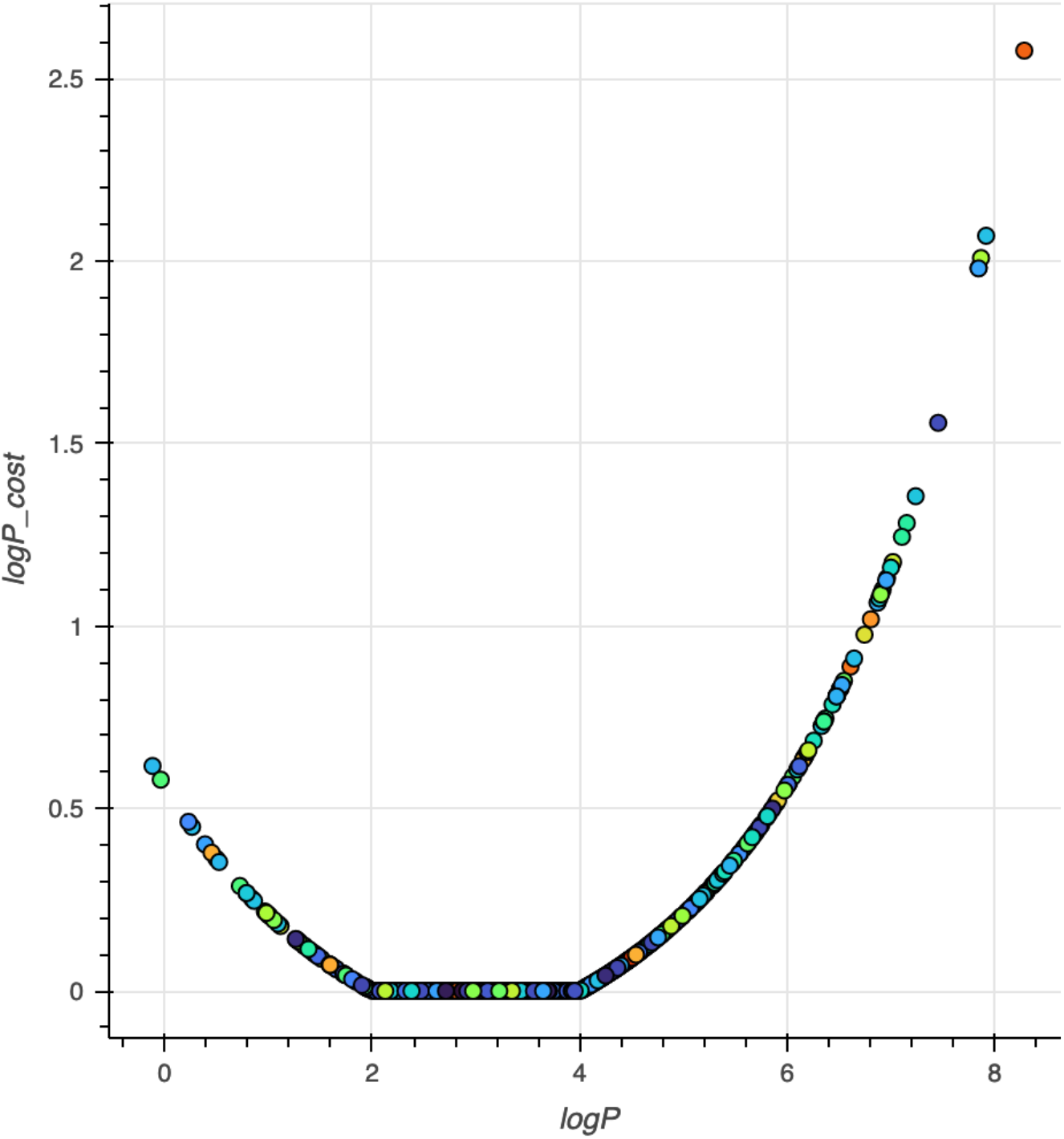
Cost term function for logP

### Detailed Predictions and Experimental Measurements for Synthesized Compounds

A table detailing the measured and predicted IC50 and Ki values for the 106 synthesized compound preparations is provided in Excel format as a supplementary data file.

